# Evaluating plant growth-defence trade-offs by modelling the interaction between primary and secondary metabolism

**DOI:** 10.1101/2024.09.15.613124

**Authors:** Jan Zrimec, Sandra Correa, Maja Zagorščak, Marko Petek, Carissa Bleker, Katja Stare, Christian Schuy, Sophia Sonnewald, Kristina Gruden, Zoran Nikoloski

## Abstract

Understanding the molecular mechanisms behind plant response to stress can enhance breeding strategies and help us design crop varieties with improved stress tolerance, yield and quality. To investigate resource redistribution from growth-to defence-related processes in an essential tuber crop, potato, here we generate a large-scale compartmentalised genome-scale metabolic model, Potato-GEM. Apart from a large-scale reconstruction of primary metabolism, the model includes the full known potato secondary metabolism, spanning over 600 reactions that facilitate the biosynthesis of 182 distinct potato secondary metabolites. Constraint-based modelling identifies that the activation of the largest amount of secondary (defence) pathways occurs at a decrease of the relative growth rate of potato leaf, due to the costs incurred by defence. We then obtain transcriptomics data from experiments exposing potato leaves to two biotic stress scenarios, a herbivore and a viral pathogen, and apply it as constraints to produce condition-specific models. We show that these models recapitulate experimentally observed decreases in relative growth rates under treatment, enabling us to pinpoint the metabolic rewiring underlying growth-defence trade-offs. Potato-GEM thus presents a useful resource to study and broaden our understanding of potato and general plant defence responses under stress conditions.

## 1. Introduction

The challenge of ensuring a secure supply of food for the rising global population is linked to improving not just the yield and quality but also the stress tolerance of major crops ^1,2^. Environmental stresses lead to annual losses amounting to billions of euros per crop. Apart from the detrimental effects of abiotic stresses, such as temperature changes, droughts and floods, biotic stresses lead to yearly losses of up to 80% of crop yield ^3–5^. In the case of potato, especially damaging are viral infections and herbivore infestations, including Potato virus Y (PVY) ^6^ and Colorado potato beetle (CPB) ^7,8^, respectively. Despite these concerns, the molecular processes underpinning and associating crop yield and defence responses are still not well understood ^9,10^. Plants attacked by biotic stressors slow down their growth to preserve molecular resources and direct them for defence purposes, including production of signalling as well as defence compounds ^9^. Conversely, rapid plant growth to improve accessibility of resources (e.g. when seeking light during germination or due to a crowded environment) is often accompanied by increased susceptibility to pests and pathogens, as growth is prioritised over defence ^11^. This growth–defence trade-off is a fundamental principle of plant economics, allowing plants to balance growth and defence according to external conditions ^9,10^. However, modern agricultural crops, including potato, have been bred to maximise yield– and growth-related traits at the expense of losing useful defence-related traits ^12^. To this end, improved understanding of the molecular mechanisms behind growth-defence trade-offs is a crucial step toward enhancing breeding strategies that could help design superior crops, combining high yields with the ability to defend against stress ^1,13^.

The stress response is often systemic in that it occurs throughout the plant and beyond the infected or damaged tissue ^14^. It is mediated by complex signalling and regulatory networks which sense and respond to environmental perturbations ^15,16^. Hormones, like salicylic acid and jasmonic acid, induce plant resistance mechanisms to either biotrophic pathogens, such as PVY ^17,18^, or herbivores, such as CPB ^7^, respectively. At the interface of growth and defence lies metabolism, which comprises a complex network of biochemical reactions that synthesise and transform substances into energy and base components necessary for the various cellular tasks ^19,20^. Here, plant growth is mediated by primary, biomass-producing processes that include photosynthesis, respiration, and the synthesis and degradation of carbohydrates, amino acids and nucleic acids ^21^, whereas secondary metabolism facilitates the production of a plethora of signalling and defence compounds (so-called specialised metabolites) ^22,23^. Lipid metabolism is also a critical subsystem interlinking the growth and defence processes ^24^, by providing precursors for many molecules in secondary metabolism and signalling pathways. Although plant metabolism has been thoroughly modelled and studied with constraint-based mathematical modelling approaches based on genome-scale metabolic models (GEMs) ^25–29^, the lack of secondary metabolism reconstructions hinders the ability of these approaches to study growth-defence trade-offs in the context of metabolism. Metabolic flux trade-offs in plants have been studied in the context of constraint-based modeling, but focused on models of primary metabolism of *Arabidopsis thaliana* ^30,31^ as well as highly condensed representation of metabolism of diverse fruits ^32^. Therefore, to correctly interpret experimental data and study the effects of plant biotic interactions at the molecular level, it is imperative to refine and expand existing metabolic modelling resources to encompass not merely the primary but also the full secondary metabolism in an essential crop system, such as potato.

In the present study, we present Potato-GEM, a metabolic reconstruction of potato leaf metabolism that spans not only all critical primary and lipid metabolic processes, but also adds a full reconstruction of the known potato secondary metabolism. We then perform a general analysis of how secondary metabolite production is linked to plant growth, determining the models’ inherent potential to capture growth-defence trade-offs and to predict how resource limitation affects these trade-offs. We further process and analyse transcriptomic data from biotic stress experiments on potato leaves, capturing both insect pests (chewing herbivore, CPB) and pathogens (intracellular virus, PVY) interaction characteristics. To connect the enzyme-catalyzed reactions of Potato-GEM with the underlying genes, we use the transcriptomic data to constrain reaction upper bounds and build a set of condition-specific models. We find that these models indeed reflect experimental observations of decreased growth under stress conditions, serving as excellent starting points for analysis of growth-defence trade-offs. Finally, we perform an in-depth analysis of the condition-specific models using Monte Carlo sampling and pathway enrichment analysis, obtaining further insights into the metabolic rewiring underpinning potato growth-defence trade-offs. Our study thus demonstrates the usefulness of secondary metabolism-expanded models, such as Potato-GEM, in the context of constraint-based modelling approaches, to help expand our knowledge and understanding of the molecular principles behind plant stress responses and environmental interactions.

## 2. Results

### 2.1 Constructing Potato-GEM by merging and curating multiple metabolic modules

To ensure an accurate reconstruction of potato metabolism, we first merged the metabolic model of *Arabidopsis thaliana* core metabolism (AraCore, spanning 549 reactions and 407 metabolites) ^25^ and the basic single-tissue model of tomato metabolism, recently updated and expanded in the Virtual Young TOmato Plant (VYTOP, spanning 2,261 reactions and 2,097 metabolites) ^27^ (Methods M1). The rationale for initiating Potato-GEM from these models is that tomato is a genetically and metabolically closely related plant from the *Solanaceae* family ^33^, whereas the Arabidopsis core metabolism constitutes the set of functionally conserved metabolic pathways across the dicot species ^25^. During the merging process, we identified an overlap of 332 reactions and 346 metabolites common to both models (Figure 1A). However, the resulting model did not contain a functional secondary metabolism nor was it able to produce vital lipid-related precursors. To resolve this issue, we further integrated the model with the recently developed Plant Lipid Module ^24^, identifying an overlap of 296 reactions and 403 metabolites present in the lipid module and the merged AraCore-VYTOP model (Figure 1A). Moreover, we curated 363 reactions from 107 pathways belonging to potato secondary metabolism and 22 related precursor pathways from the MetaCyc-derived Plant Metabolic Network database ^34,35^, adding an additional 277 reactions and 256 metabolites (Methods M1). This resulted in the Potato-GEM model spanning 8,272 reactions and 4,982 metabolites (Figure 1A), across 16 unique compartments (Figure 1B: only key compartments shown, since the majority are related to the Plant Lipid Module, Supp. files S1, S2). Compared with the tomato GEM in VYTOP, the number of blocked reactions (i.e., those unable to transport any flux) was reduced ∼5-fold (Figure 1C: ratio of blocked reactions was 10.5% with Potato-GEM and 51.4% with VYTOP).

**Figure 1.**
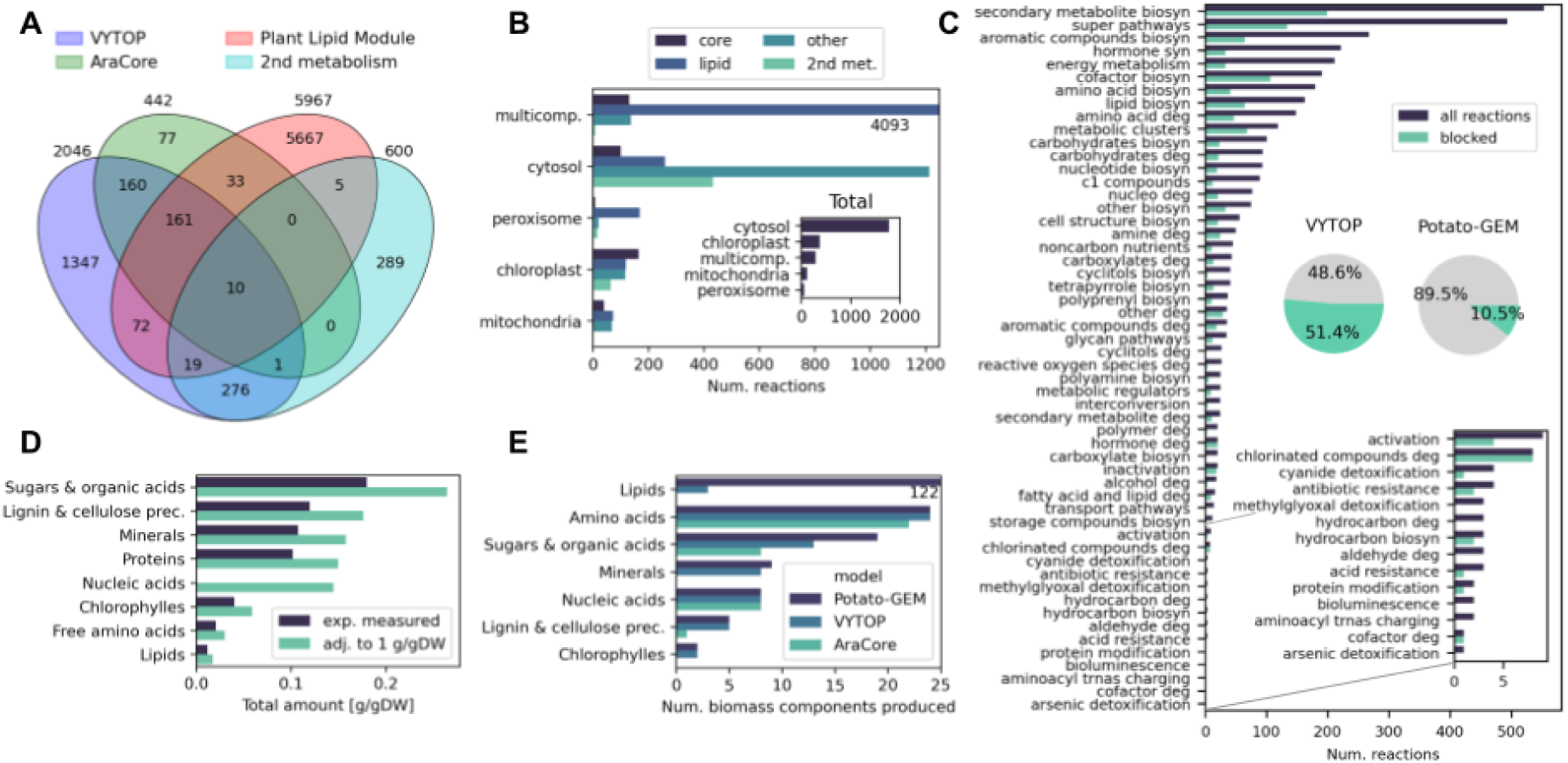
Constructing Potato-GEM by merging and curating multiple modules. (A) Venn diagram of the four models that comprise potato-GEM, including AraCore ^25^, the basic single-tissue model from Virtual Young TOmato Plant (VYTOP) ^27^, Plant Lipid Module ^24^ and additional manual curation of secondary metabolism based on the MetaCyc database ^35,45^. (B) Number of core, lipid and other primary as well as secondary metabolism reactions across the model’s key cellular compartments including multi-compartment reactions. Inset shows the total number of reactions. (C) Depiction of the total number and blocked reactions across the MetaCyc pathway ontology. Inset pie charts show the ratio of blocked reactions in VYTOP and potato-GEM. Lower inset shows a zoom in on the values of the bottom 14 pathways. (D) Depiction of potato biomass components based on experimentally measured values and after adjustment of total content to 1 g/gDW ^43^. (E) Quantification of the capability to produce biomass precursors across potato-GEM and the existing VYTOP ^27^ and AraCore ^25^ models.

We next defined a leaf biomass growth reaction to use with Potato-GEM metabolic simulations. Here, we measured the dry weight to fresh weight ratio as well as the total protein content of potato leaves (Methods M3). Additionally, quantitative data for various biomass components, including sugars, organic acids, amino acids, and lipids, were compiled through an extensive review of the literature ^24,27,36–42^. The leaf biomass thus comprised 67 compounds and an additional 122 lipid backbones and chains (Figure 1D, Supp. table S1, Supp. file S3). To ensure correct optimization, the component values were further calibrated to a total of 1 g/gDW ^43^ by proportionally increasing their quantities and setting the overall quantity of nucleic acids, for which experimental data was not available, similar to that of proteins (Figure 1D: 14.5% of DW) ^27^. We then tested if Potato-GEM can be used to simulate growth under phototrophic (light) and heterotrophic (dark) nutrient-limiting conditions (Methods M2). Under both light and dark regimes, we found that the model is indeed capable of producing all 189 biomass components (Supp. Figure S1.1). Moreover, compared with the existing AraCore and VYTOP models, Potato-GEM includes a larger number of biomass precursors across multiple component classes (Figure 1E: producing in total 71.8% or 11.7% more biomass precursors if lipids are disregarded, respectively), supporting its secondary metabolite-producing functionality.

Importantly, the reconstruction of secondary metabolism in Potato-GEM captures the complete *Solanum tuberosum* secondary metabolism as detailed in the Plant Metabolic Network database ^34,35^ (Methods M1). It covers the major classes including: (i) alkaloids, such as: alpha-solanine and alpha-chaconine, calystegines and tropane alkaloids, (ii) phenylpropanoid derivatives, such as: flavonoids, coumarins, cinnamates, lignans and lignins, (iii) terpenoids, incl. carotenoids and mono-, di-, tri– and sesquiterpenoids, (iv) phytoalexins, i.e. resveratrol and capsidiol, and (v) hormones, including jasmonic acid, salicylic acid, abscisic acid, auxins, brassinosteroids, cytokinins, ethylene and gibberellins (Figure 2A, Supp. figure S1.2). This enables the modelling of the production of defence compounds related to stress response including hormonal biosynthesis, allowing us to study the effects and coupling of growth with secondary metabolite production ^19,44^. The complete module spans over 600 reactions in 107 pathways, producing a total of 182 unique secondary metabolites (Supp. figures S1.2, S1.3, Supp. table S2, Supp. file S4).

**Figure 2.**
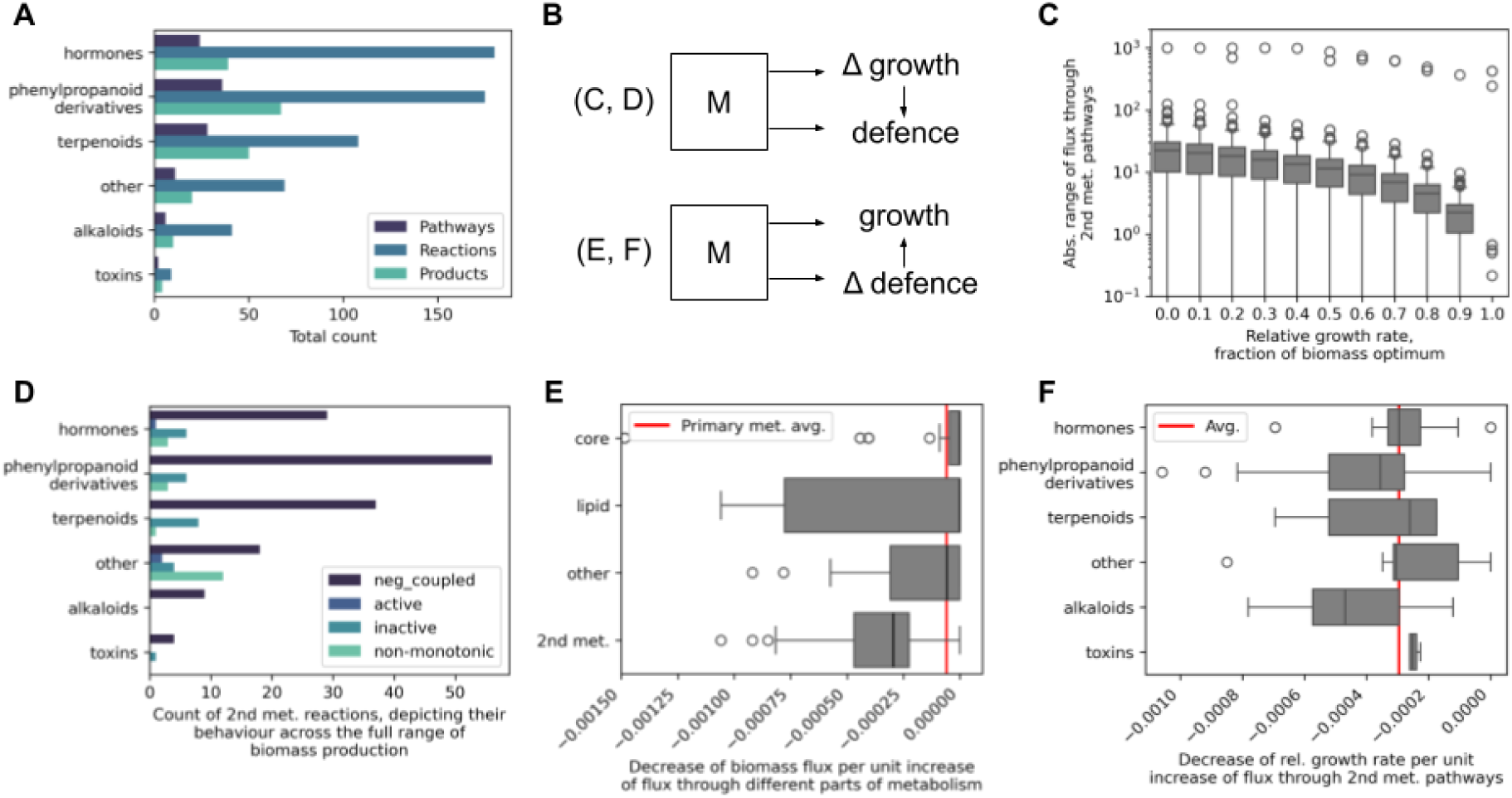
Secondary metabolism reconstruction enables quantifying growth-defence trade-offs. (A) The number of reactions, pathways and products across secondary metabolism classes in the reconstruction. (B) Schematic depiction of the studied growth-defence trade-off relationships by either observing the effects of growth limitation on defence activation (as depicted in panels C, D) or vice versa (panels E, F). (C) Depiction of predicted growth-defence trade-off capabilities as the absolute range of flux through secondary metabolic pathways in response to the varying ratio of optimal biomass production in FVA. (D) Segregation of secondary pathway reactions across their coupling behaviour with the growth objective, showing that the majority of these reactions are negatively coupled with growth and exhibit growth-defence trade-offs. (E) Distribution of the decrease of biomass flux per unit increase of flux through different parts of metabolism including core, lipid and secondary metabolism. Red line denotes the median value of primary metabolism. (F) Distribution of the decrease of biomass flux per unit increase of flux through pathways across secondary metabolism classes. Red line denotes median value across the whole secondary metabolism.

### 2.2 Secondary metabolism reconstruction enables quantifying growth-defence trade-offs

The refined Potato-GEM model enabled us to investigate the coupling ^46^ between potato growth and stress response modes and to ascertain the possibility of growth-defence trade-offs. To this end, we investigated if fluxes through secondary pathways and reactions are coupled to biomass production (growth, Figure 2B). Here, the variability of fluxes of the final product-producing reactions in each secondary pathway were evaluated within the optimal biomass space using flux variability analysis (FVA) ^47^, where different fractions, ranging from 0 to 1, of the optimal relative growth rate were used (Figure 2C). We observed that at the optimal relative growth rate, the majority (96%) of secondary pathways were either inactive, with computed flux ranges equalling 0 h^-1^, or minimally active, with a limited flux range below 1 h^-1^, as occurred with 6 reactions (Figure 2C, Supp. figure S2.1). The exception was cytokinin biosynthesis (PWY-2681) that was positively coupled with growth, as its flux lower bound increased proportionally to the relative growth rate (Supp. figure S2.2). Furthermore, with a decreasing relative growth rate from the optimum, the majority of secondary pathways exhibited proportional increases of their flux ranges (Figure 2C), supported by a significant negative correlation (Spearman *ρ* = –0.55, *p*-value < 10^-16^, Supp. figure S2.3) between growth and flux through secondary metabolite production. This demonstrated that the majority of secondary pathways are thus negatively coupled with the plants’ growth objective (Figure 2D: 155 reactions across 86 pathways). In addition, of these negatively coupled pathways, almost all (98%) were found to be strongly negatively correlated (Spearman *ρ* < –0.99, *p*-value < 8×10^-12^, Supp. figure S2.3) with the relative growth rate. The remaining pathways were either positively coupled (1 reaction and pathway), exhibited non-monotonic and thus unstable changes (17 reactions across 9 pathways), or were inactive across the whole range of biomass production (25 reactions across 20 pathways, Figure 2D). Importantly, an optimal secondary metabolite production was observed at a relative growth rate of 0.4, where the largest amount of secondary reactions and pathways were found to be active (Supp. figure S2.1).

The observed negative coupling between growth and secondary metabolism (Figure 2C) suggested that a growth-defence trade-off is occurring with these pathways. Therefore, we further investigated how much a unit increase of flux through secondary pathway production (defence) decreases the flux through the biomass reaction (growth), which can be interpreted as a growth-defence trade-off factor (Figure 2B, Methods M2). On average, across all coupled secondary pathways (Figure 2D), we observed a 3×10^-4^ decrease of biomass flux with an unit increase of secondary metabolite production (Figure 2E). Interestingly, this growth-defence trade-off factor was ∼5-fold and significantly (Wilcoxon rank-sum test *p*-value < 0.0052) higher than a trade-off factor obtained in response to unit increase of flux through primary metabolic reactions (Figure 2E: ∼10^-5^). With primary metabolic response, both the core and lipid metabolism resulted in the lowest trade-offs on average (Figure 2E). Moreover, we observed that the growth-defence trade-off factors remain on average relatively equal across different secondary metabolite classes (Figure 2F). Here, the largest factor of 4.7×10^-4^, ∼1.6-fold larger than the overall secondary metabolism average, was found for alkaloids, likely due to the large costs for their synthesis^48^. Overall, the results suggest that under stress, secondary metabolism activation under defence responses requires a relatively higher amount of resources (Figure 2E: > 5-fold more) than the general rewiring of core metabolism.

### 2.3 Exploring the effects of resource limitation on growth and defence

We next explored the hypothesis that resource constraint is a primary reason for the inverse growth-defence relationship, meaning that providing more resources would reduce growth-defence trade-offs and allow plants to simultaneously grow and defend themselves ^10,49^. To this end, we proportionally limited or increased the availability of key model resource inputs: CO_2_, light, nitrogen or a combination of all three (Figure 3A, Methods M2). We observed proportional decreases in the predicted growth rates when limiting resources, reaching no growth when resources were completely withdrawn (Figure 3B: note that a resource ratio of 1 is used to denote resource consumption at the optimal relative growth rate). Conversely, as expected, no change in growth was observed when increasing resource availability above a resource ratio of 1 (Supp. figure S3.1). Similarly as with growth, the secondary metabolite production capacity decreased significantly (Wilcoxon rank-sum test *p*-value < 10^-14^, measured between the resource ratio of 1 and 0) and in proportion to the limited resources (Figure 3C). The exception was with nitrogen limitation, where the defence response decreased by merely 6% when varying nitrogen availability between a resource ratio of 1 and 0. Increasing resource availability, on the other hand, did not lead to an increased secondary production capacity (Supp. figure S3.2), likely since the model’s optimal secondary metabolite production was already reached at the relative growth rate of 0.4 within the given constraints. The results suggest that under actual *in situ* conditions, where plants do not grow at the metabolic optimum (as depicted in Figure 3C), access to more resources could indeed decrease growth-defence trade-offs by allowing plants to simultaneously grow and defend themselves ^10^.

**Figure 3.**
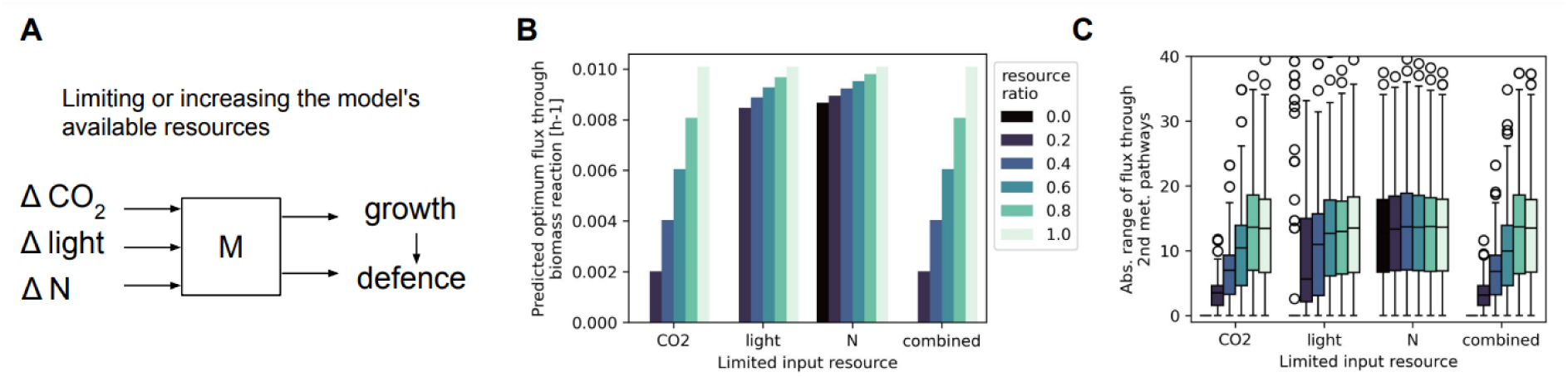
Exploring the effects of resource limitation on growth and defence. (A) Schematic depiction of the analysis of resource limitation or expansion, where the key resources CO2, light and nitrogen were varied within a range of 0 to 1 relative to the resource consumption flux obtained at an optimal growth rate of 1. (B) Effect of resource limitation on the predicted relative growth rate (biomass optimum). (C) Effect of resource limitation on the range of secondary metabolite production (defence), computed at the fraction of 0.4 of the relative growth rate to observe the largest amount of possible active secondary metabolism.

### 2.4 Capturing biotic stress responses with transcriptomics data

Our next aim was to investigate a range of common potato biotic stress scenarios and the growth-defence trade-offs that they elicit. To this end, we performed a transcriptomics experiment based on the herbivore attack of potato leaves with the Colorado potato beetle (CPB) ^50^ and complemented it with previously published data on the pathogen interaction with the potato virus Y (PVY) ^51^ (Figure 4A). Briefly, within the presently performed CPB experiment, plants were exposed to two beetles per leaf for 30 minutes and 24 hours post infection, the leaf region surrounding the damaged part was sampled in parallel with non-infested leaves (control) and processed for RNA-Seq (Methods M3,4). In the previously published PVY experiment ^18^, leaves were inoculated with PVY and tissue immediately surrounding the site of viral multiplication was sampled when the hypersensitive resistance response was fully established (4 days after inoculation), in parallel with control non-inoculated leaves tissue, and processed for RNA-Seq (Methods M4). In both experiments, biological triplicates were analysed per treatment.

**Figure 4.**
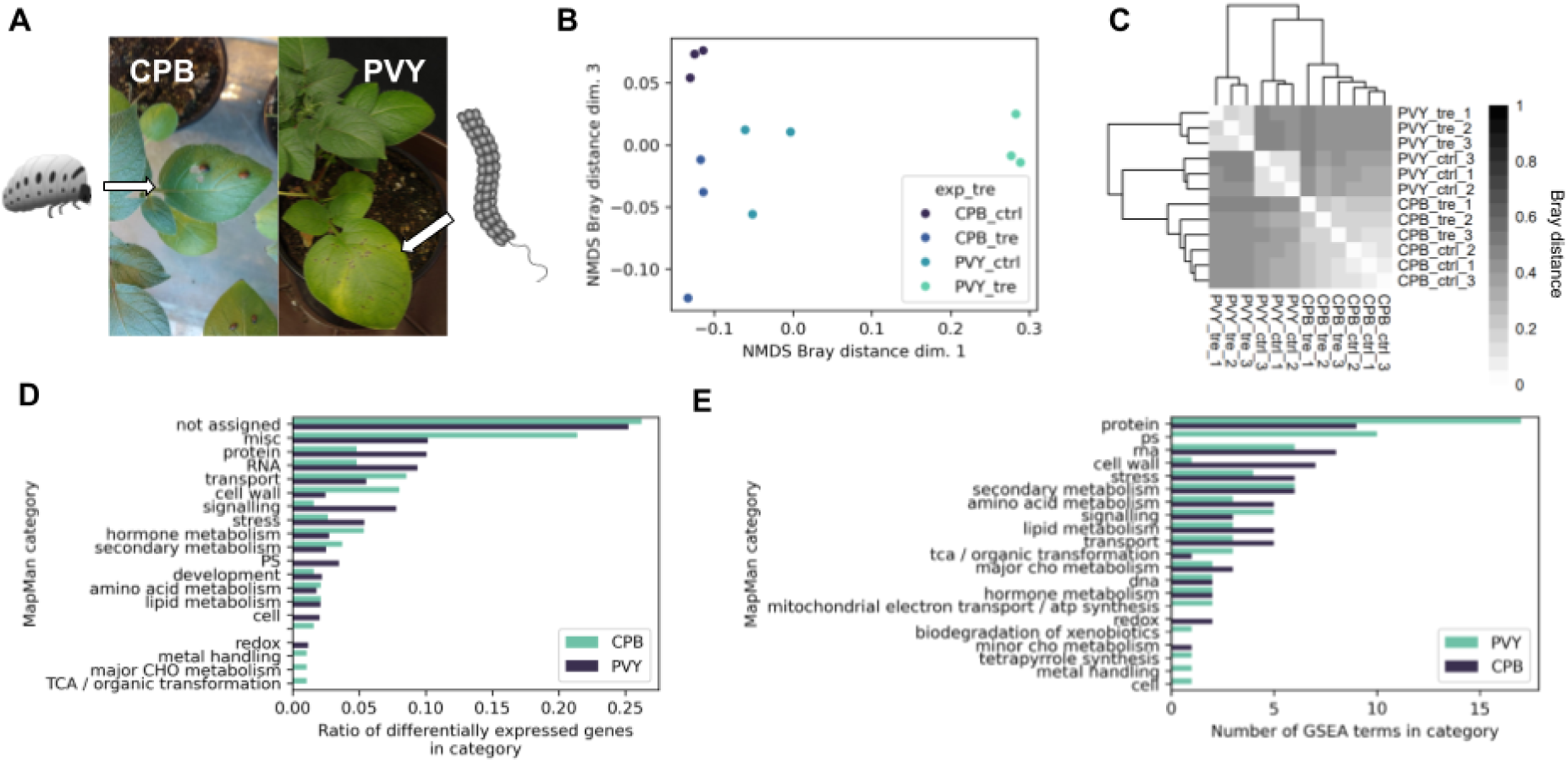
Capturing biotic stress responses with transcriptomics. (A) Schematic depiction of biotic stress transcriptomics experiments with Colorado potato beetle (CPB) and Potato virus Y (PVY). (B) Principal-coordinate analysis of Bray-Curtis dissimilarity between transcript counts across experiments and treatments. (C) Heatmap and dendrogram visualisation of hierarchical clustering on Bray-Curtis dissimilarity between sample replicates across experiments and treatments. (D) Ratio of differentially expressed genes (DEGs) (BH-corrected *p*-value < 0.05 and abs(log2FC) > 2) belonging to a MapMan ^52,59^ category versus the total number of DEGs across experiments. Ratios above 0.025 shown. (E) Transcriptomics responses on the level of pathways. MapMan-defined pathways ^52,59^ enriched with differentially regulated genes were calculated using GSEA ^60^ (BH-corrected *p*-value < 0.05).

Non-metric multidimensional scaling (NMDS) analysis of transcript counts indicated clear separation among treated and control samples in both experiments (Figure 4B, Supp. figure S4.1, Methods M4). We also observed significant correlation (Spearman *ρ* = 0.775, *p*-value < 10^-16^) of transcript counts among the control samples of both experiments (Supp. Figure S4.2). However, whereas correlation analysis among all 12 samples showed a clear distinction between PVY control and treatment samples, this was not observed with CPB samples (Figure 4C). The CPB experiment also captured a smaller number of differentially expressed genes (325 DEGs), compared to the PVY experiment (9,289 DEGs) (Figure 4D, Supp. file S5, see Methods M4). These differences were likely due to the shorter CPB exposure time and thus milder observed defence response than with PVY ^18^. Nevertheless, further DEG and enrichment analysis using plant specific MapMan ontology terms ^52^ showed (i) an increase in hormonal production, specifically salicylic acid with PVY ^17^ and jasmonic acid with CPB ^8,53^, which are known signalling cascade regulators of the specific biotic stresses, (ii) general upregulation of multiple secondary pathways, including those belonging to glycoalkaloid ^54^, phenylpropanoid ^55,56^ and terpenoid classes ^57^, (iii) lowered photosynthetic activity with PVY ^58^, (iv) upregulated sucrose degradation factors with PVY likely due to increased sucrose accumulation ^56^, as well as (v) downregulation of biotic stress response-related factors due to an overactivated signalling response (Figure 4E, Supp. file S6) ^16^. This suggested that known responses as a consequence of biotic stress and growth-defence trade-offs were indeed captured within both experiments ^18^.

### 2.5 Transcriptome-constrained models recapitulate reduced growth in stress response

To further study the characteristics of biotic interactions, we next constructed models constrained by the transcriptomics data (Figure 5A). To this end, we first annotated model reactions with gene protein reaction (GPR) associations, which were obtained from the MetaCyc-derived Plant Metabolic Network database ^34^ (*Solanum tuberosum* subset), as well as by translating GPRs with Arabidopsis gene identifiers from the Plant Lipid Module ^24^ and AraCore ^25^ modules (Methods M5). Here, orthologous genes between Arabidopsis and potato were identified according to Plaza v5.0 ^61^ orthologs using clustering algorithms as well as BLAST reciprocal best hit search ^62,63^. This yielded a set of 3,134 potato gene identifiers and resulted in 3,950 reactions (48%) annotated with GPR associations comprising, on average, 2 unique gene identifiers (Figure 5B). Unsurprisingly, better coverage of GPR-annotated reactions of 65% was achieved with the core metabolism compared to a 50% coverage of both the lipid and secondary metabolism (Figure 5B inset). However, secondary metabolic reactions were annotated with GPR associations that comprised significantly (Wilcoxon rank-sum test *p*-value < 3.3×10^-13^) more unique gene identifiers than primary metabolic ones, with a 1.5-fold higher number of genes on average (Figure 5C). Within the secondary metabolism, apart from precursors exhibiting the highest coverage (83%), terpenoid, toxin and hormone classes achieved a better than average coverage of over 48% (Figure 5D).

**Figure 5.**
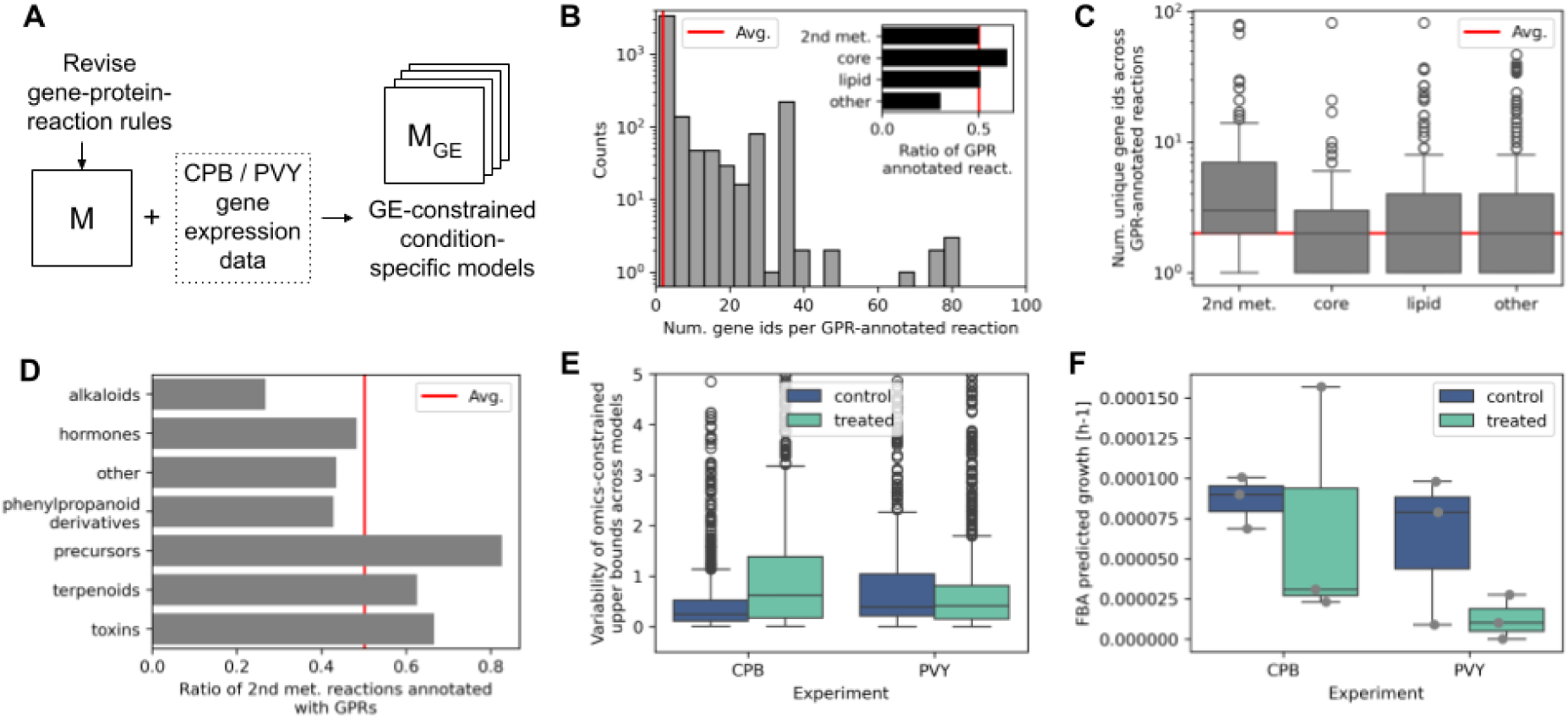
Transcriptome-constrained models recapitulate reduced growth in stress response. (A) Schematic depiction of the procedure to construct a set of CPB– and PVY-transcriptome constrained models. (B) Number of gene identifiers present in GPR associations across GPR-annotated reactions. Red line denotes an average value of 2. Inset depicts the ratio of GPR-annotated reactions across the general metabolic types. (C) Number of gene identifiers present in GPR associations of GPR-annotated reactions across the general metabolic types. Red line denotes the average value. (D) Ratio of GPR-annotated reactions across secondary metabolism classes. Red line denotes the average value. (E) Variability of reaction upper bounds across replicates in the CPB– and PVY-transcriptome constrained models. (F) Predicted biomass production with the transcriptome constrained models under a phototrophic regime.

Next, we integrated the transcriptomics data with the Potato-GEM model, by evaluating GPR associations based on the provided transcriptomic measurements to constrain the model’s upper bounds of flux across reactions. This resulted in 12 transcriptomics-constrained models, one for each replicate of the data treatment (control and treated) and experiment type (CPB and PVY, see Figure 4). An initial 14,404 and 19,367 transcripts from the CPB or PVY experiment, respectively, were mapped to GPR associations across 3,561 and 3,467 reactions, respectively (Supp. figure S5.1). Of the GPR-annotated reactions, ∼46% (Supp. figure S5.1: 1,725 and 1,692, respectively) were essential, without which the model cannot produce the target biomass (i.e. no growth) ^64^. We observed that the variability of the upper bounds of the CPB constrained models differed significantly (Wilcoxon rank-sum test *p*-value < 10^-16^) across treatments, where models corresponding to treated samples exhibited an average 2.5-fold increase in variability of upper bounds compared to controls (Figure 5E). However, this was not the case with PVY, where the variability remained approximately equal between treated and control models. Similarly, no significant difference in upper bound variability was observed for essential in comparison to non-essential reactions (Supp. figure S5.2). Moreover, the transcriptomics-constrained reactions were found to comprise the full range of pathways covering 96% of the key cellular processes as existing in the original unconstrained model (Supp. figure S5.3, see Figure 1C).

Finally, we observed that under both biotic stress scenarios, the predicted relative growth rates (Methods M2: biomass production under phototrophic regime as applied in the experimental setup) with the constrained models of treated plants were on average 73% lower than that of controls (Figure 5F: 0.39 of the optimal growth rate observed with CPB and 0.14 with PVY, respectively, note that significance not observed due to low amount of replicates). This is in line with experimental observations, where growth rates under biotic stress treatments have generally been found to be lower compared to controls ^65,66^. This suggested that the transcriptomics-constrained models represent accurate metabolic replicas of the *in vivo* stress scenarios, supporting their usefulness to further study biotic stress-induced growth-defence trade-offs in the context of metabolism.

### 2.6 Exploring the mechanisms of growth-defence trade-offs under biotic stress

We next set out to perform a large-scale treatment-specific analysis of metabolism to obtain a global picture of metabolic responses under biotic stress. To this end, we applied randomised Monte Carlo sampling using the Artificial centred hit and run (ACHR) algorithm^67^ (Figure 6A, Methods M2). To ensure that growth-defence trade-offs were appropriately captured in the sampling procedure, the global reaction bounds used for sampling were obtained from FVA performed over a range of fractions of the biomass optimum, between optimal secondary metabolism production (at the fraction of 0.4) and optimal biomass production (fraction of 1, see Figure 2C, Methods M2). Additionally, we were interested in observing metabolic states related to known hormone responses that are mediated by extensive signalling cascades ^15^, which are not the part of the present metabolic model. We thus imposed that the transcriptomics-constrained models of non-stressed control conditions produce no jasmonic acid with CPB and no salicylic acid with PVY, respectively, based on experimental and published observations ^8,17,53^ (Methods M2).

**Figure 6.**
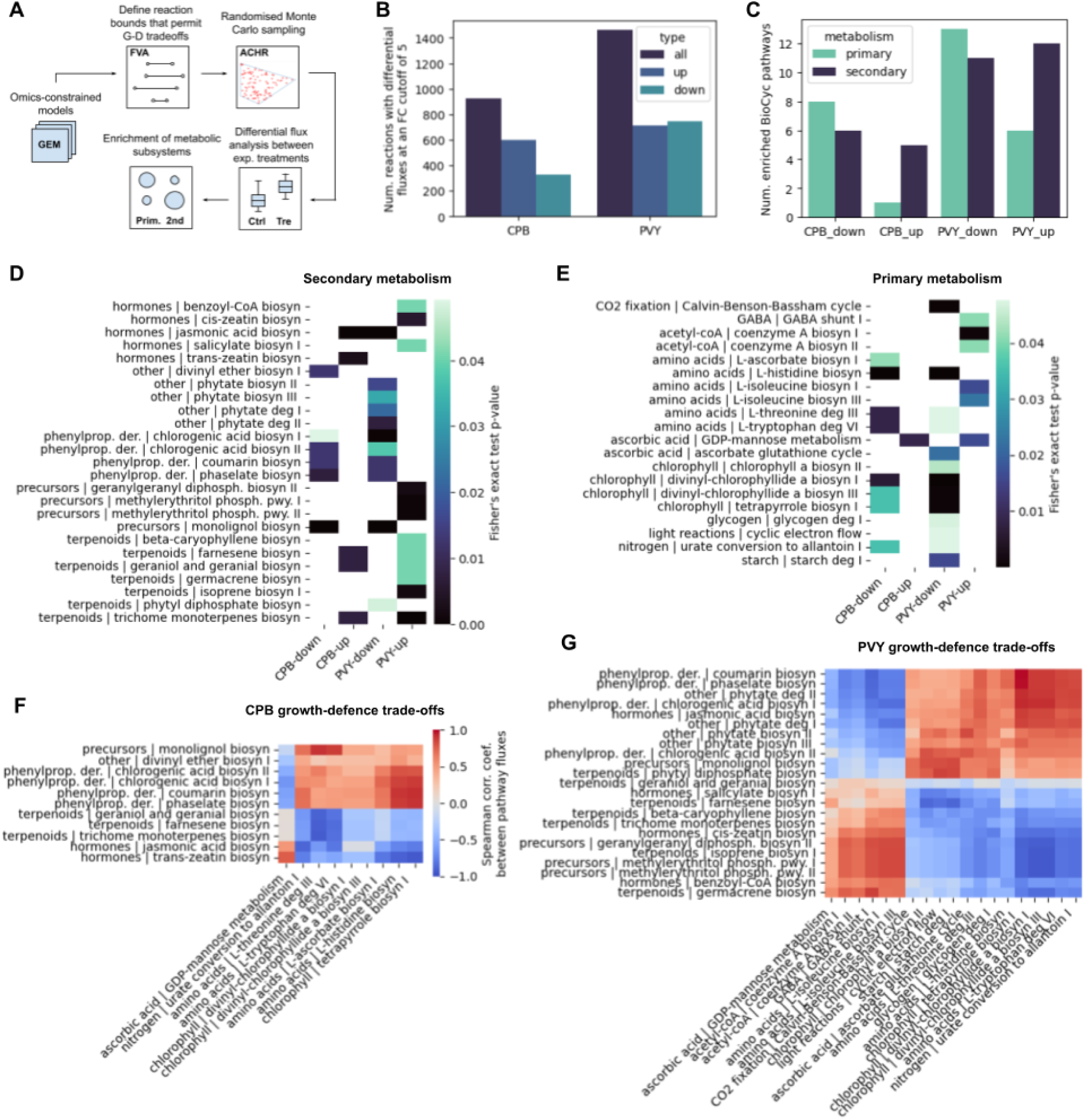
Exploring the mechanisms of growth-defence trade-offs under biotic stress. (A) Schematic depiction of the randomised sampling procedure applied to investigate the enrichment of metabolic subsystems among biotic stress treatment and control models. (B) Number of identified reactions with differential fluxes at a fold-change cutoff of 5 (Kolmogorov Smirnov test BH-corrected *p*-value < 0.05). Upregulated (up), downregulated (down) and total (all) number of reactions depicted separately. (C) The number of significantly (Fisher’s exact test BH-corrected *p*-value < 0.05) up– and down-regulated BioCyc pathways according to differential reactions. (D) Heatmap of secondary metabolism BioCyc pathways significantly (Fisher’s exact test BH-corrected *p*-value < 0.05) enriched in differential reactions. Class and full pathway names provided in Supp. Table S4. (E) Heatmap of primary metabolism BioCyc pathways significantly (Fisher’s exact test BH-corrected *p*-value < 0.05) enriched in differential reactions. (F,G) Correlations between total fluxes of the enriched (Fisher’s exact test BH-corrected *p*-value < 0.05) secondary vs. primary pathways, demonstrating growth-defence tradeoffs, with (F) CPB and (G) PVY models. Dendrograms depicting grouping of profiles shown in Supp. figure S6.5. Cosine distance metric used and Spearman correlation coefficient shown (*n* = 1,000 samples each).

We then performed differential flux analysis to identify reactions that exhibited significant (Kolmogorov Smirnov test BH-corrected *p*-value < 0.05) and over two-fold differences in flux values between treatment and control conditions, on average (Methods M6, note that replicates were pooled for these computations). In relation to the fold change cutoff used, which was varied between two– and ten-fold, the analysis revealed between 1,958 and 601 (24% and 7%) differential reactions with CPB, and between 2,519 and 1,012 with PVY (30% and 12%), respectively (Figure 6B, Supp. figure S6.1). With PVY, we observed ∼1.5-fold more differential reactions than with CPB, irrespective of the fold change cutoff (Supp. figure 6.1). With both experiments, the fluxes of differential reactions were on average ∼40% and significantly (Wilcoxon rank-sum test *p*-value < 0.014) lower in treatment models compared to control ones (Supp. figure S6.2). This was likely due to the 82% higher number of downregulated reactions compared to up-regulated ones observed with CPB, though with PVY, the number of up– and downregulated reactions was approximately equal (Figure 6B).

To determine the pathways comprising the reactions exhibiting differential fluxes across conditions, we next performed enrichment analysis of metabolic subsystems at the level of Biocyc ontologies (Supp. figure S6.3, Supp. table S3) and Biocyc pathways (Figure 6C-E, Supp. table S4). We found that CPB models were significantly (Fisher’s exact test BH-corrected p-value < 0.05) enriched in 20 BioCyc pathways across 9 ontologies, whereas PVY models spanned 43 pathways across 16 ontologies (Figure 6D,E). In both experiments, multiple secondary metabolism pathways displayed differential fluxes (Figure 6C: a total of 11 pathways with CPB and 24 with PVY, respectively), with an approximately equal number of secondary pathways exhibiting down-or upregulation. Interestingly, we observed that different secondary pathways were regulated under the different biotic stresses (Figure 6D). For instance, viral infection caused the induction of hormonal pathways, including salicylic acid and cis-zeatin, as well as multiple terpenoids including their precursors, while insect feeding led to increased synthesis of jasmonic acid and trans-zeatin hormones as well as terpenoid pathways. With both biotic stresses, multiple phenylpropanoid pathways were downregulated. Indeed, besides salicylic and jasmonic acid, both cytokinins are known regulators of plant stress responses, with a role in leaf development ^68,69^, whereas terpenoids have been found to protect plants from both viral infections and herbivores ^70,71^, with terpenoid volatiles, such as farnesene, beta-caryophyllene and germacrene, inducing indirect defences and priming neighbouring plants ^72^. On the other hand, the primary metabolism was highly downregulated in both CPB and PVY treatment models, with over two-fold fewer upregulated than downregulated pathways (Figure 6C,E). Here, with both biotic stresses, chlorophyll synthesis was significantly downregulated, in line with the expected rewiring of photosynthesis mediated by the biotic stress ^58,73^.

Next, we analysed the correlation between the sampled fluxes of the enriched secondary and primary pathways under the biotic stress conditions (Figure 6F,G, Supp. figure S6.4). Fluxes were summed over the respective reactions per pathway to obtain the total pathway flux (Methods M6). Indeed, significant correlations (|Spearman corr. coef.| > 0.237, p-value < 10^-16^) were found among almost all pathways (Figure 6F,G: secondary vs. primary metabolism pathways shown depicting growth-defence trade-offs). We further observed that specific groups of pathways demonstrated similar growth-defence trade-off patterns in the context of secondary vs. primary pathway flux correlations (Supp. figure S6.5). By calculating the cosine distance and performing permutational tests, 4 significantly distinct clusters (permutation test p-value < 0.01) were identified with the CPB models (Figure 6F, Supp. figure S6.5A, Methods M6). Interestingly, here we observed a cluster showing negative correlation (Spearman *ρ* < –0.81, *p*-value < 10^-16^) between fluxes in terpenoid synthesis and amino acids Thr and Trp degradation, as well as a separate cluster linking decreased chlorophyll and histidine synthesis with phenylpropanoid synthesis (Spearman *ρ* > 0.35, *p*-value < 10^-16^). On the other hand, with PVY, 5 significantly distinct clusters (permutation test *p*-value < 0.01) were observed (Figure 6G, Supp. figure S6.5B). Similarly as with CPB, fluxes in terpenoid synthesis as well as the hormone cis-zeatin were negatively correlated (Spearman *ρ* < –0.45, *p*-value < 10^-16^) with fluxes in Thr and Trp degradation and additionally His synthesis. On the other hand, downregulated phenylpropanoid derivative production negatively correlated (Spearman *ρ* < –0.43, *p*-value < 10^-16^) with the synthesis of Ile and GABA, and positively (Spearman *ρ* > 0.39, *p*-value < 10^-16^) with decreased chlorophyll synthesis. This showcases how the specific primary metabolic rewiring, which underlies secondary defence activation and production under stress conditions, can be pinpointed, and suggests the existence of different metabolic trade-off “programmes” useful for further study.

## 3. Discussion

In the present study, we asked whether we can explain plant growth-defence trade-offs through metabolic flux signatures obtained by constructing and analyzing a sufficiently comprehensive and accurate model of primary and secondary metabolism in the major crop plant, potato. To this end, we developed and applied potato-GEM, a genome-scale metabolic model constructed by merging multiple models ^25,27^ and modules ^24^ (Figure 1A), and includes a reconstruction of the full potato secondary metabolism according to the Plant metabolic network ^34^ MetaCyc database ^35^ (Figure 2A). We demonstrated that through its comprehensiveness and the secondary metabolism reconstruction, the model enables exploring the principles of plant growth-defence trade-offs in the context of metabolism and under biotic stress.

Similarly to growth ^25,27^, plant responses to abiotic and biotic stress comprise a large set of cellular processes ^15,16,44^. In contrast to targeted *in planta* experimental research that may be limited to studying merely a subset of different molecular responses at once ^9^, *in silico* mathematical modelling of cellular processes such as metabolism ^24,25^ and signalling ^15,16^ enables us to jointly capture and examine the full range of cellular responses. The potato-GEM model thus allowed us to observe, explore and interpret the rewiring of plant metabolism from growth to defence-driven responses. The analysis of growth-defence trade-offs was performed in two ways. First, we directly probed the model using flux balance and flux variability analysis (FBA and FVA, respectively) ^47,74^ (Figure 2B and 3A), either by (i) decreasing the relative growth rate (fraction of biomass optimum) and observing how defence pathways are unlocked (Figure 2C,D), (ii) varying the secondary metabolite production and determining its effects on growth (Figure 2E,F), or (iii) testing the effect of limiting different key input resources on the growth-defence trade-offs ^10,49^ (Figure 3B,C). Second, we constrained the model according to experimental gene expression measurements under biotic stress ^18^ (Figure 4) and then analysed model predictions by performing (i) FBA (Figure 5F) as well as (ii) a sequential combination of FVA and randomised sampling followed by differential flux analysis and pathway enrichment analysis ^67,75^ (Figure 6).

As a result of the first approach, we found that the effects of decreasing the relative growth rate (flux through the biomass reaction) were significantly negatively correlated with increasing flux through secondary pathways (Figure 2C), enabling us to quantify specific general principles of growth-defence trade-offs. Specifically, we identified that the largest number of defence-related secondary metabolism pathways are active at a decrease of ∼60% of the potato relative growth rate (Figure 2C: fraction of biomass optimum of 0.4). Further, compared to primary metabolism, we found that the secondary metabolism response involves more costly metabolic processes, based on measuring the effect of perturbing secondary metabolite production (Figure 2E: unit change flux through secondary pathways) on growth (flux through biomass reaction). The cost of diverted resources is also, on average, approximately equal across the secondary metabolite classes (Figure 2F). Indeed, it is known that due to the costliness of diverting energy and resources away from key processes, such as growth and reproduction ^76^, cells employ multiple strategies to lower the high metabolic costs of defence ^23^. Apart from fine-tuning secondary metabolite production via gene expression, metabolite multifunctionality and effective recycling ^23^, balancing the objectives between biomass or defence production is a key cellular strategy defined by the spectrum of resistance and tolerance of the underlying plant genotype ^76^. Lastly, we found that resource limitation has a profound effect on growth-defence trade-offs, effectively decreasing defence rewiring due to a lack of resources (Figure 3C). This supports the existing resource allocation hypothesis ^49,77^, whereby resource constraints are a primary reason for the inverse growth-defence relationship. Namely, since plants typically do not optimise merely the growth objective and thus do not necessarily grow at the metabolic optimum, they can find themselves somewhere between the minimum and maximum modelled growth rate (Figure 3B: 0 to 1 relative growth rate). Thus, providing more resources, such as increasing nutrients, CO2 and/or light (Figure 3C), would reduce growth-defence trade-offs, resulting in more secondary metabolism activation at a higher growth rate and thus allowing plants to simultaneously grow and defend ^10^.

To apply Potato-GEM to study growth-defence trade-offs under biotic stress, we first performed and processed transcriptomics experiments exposing potato leaves to CPB (this study) and PVY ^18^, respectively (Figure 4A). According to differential gene expression and pathway enrichment analyses of the transcriptomics data, both experiments captured the full expected response of their corresponding stresses (Figure 4D,E), despite the CPB experiment characterising a more early stage of the stress response than was the case with PVY ^18^. By revising the Potato-GEM gene-protein-reaction (GPR) associations that link gene expression and reactions (Figure 5A,B), we constrained reaction upper bounds according to the measured gene expression levels ^78,79^. This resulted in a set of condition-specific models, which, when predicting optimal growth rates using FBA, predicted decreased growth under biotic stress compared to controls (Figure 5F), in accordance with published experimental observations. For instance, in a study measuring the effect of CPB damage on potato growth rates in different plant growth stages, damage during both the vegetative and tuber-bulking phases led to decreased haulm and tuber growth rates ^66^. Similarly, PVY infected plants, grown from infected seed tubers, displayed slower growth rates and lower tuber yields compared to plants grown from non-infected tubers ^65^. Furthermore, especially with CPB, the ∼60% decrease in growth rate corresponded well with our findings of optimal growth-defence trade-offs at a similar decrease in relative growth rate (Figure 2C).

Finally, using a randomised Monte Carlo sampling procedure that involved differential flux and pathway enrichment analyses with the condition-specific models (Figure 6A), we observed a large and significant fraction of reactions exhibiting differential fluxes between controls and biotic-stress treatments with both experiments (Figure 6B,C). These differential flux reactions were enriched both in primary metabolic pathways as well as those involved in secondary metabolite and hormone production (Figure 6D,E), in line with previous observations ^8,80^. Here, the observed size and scope of the defence response with the CPB leaf attack was much smaller compared to the hypersensitive resistance response close to the PVY leaf entry site (Figure 6C: approx. half less pathways enriched, respectively), likely due to a shorter herbivore exposure time of 30 minutes compared to the fully developed hypersensitive response to the viral infection ^18^. Furthermore, analysis of the correlations among the sampled secondary and primary pathway flux states enabled us to directly investigate metabolic growth-defence trade-offs and how different defence pathways are activated or deactivated in accordance with specific primary metabolic changes under biotic stress (Figure 6F,G). These concerted changes likely reflect the existence of different metabolic trade-off “programmes”, and point to interesting areas of further study as they enable the development of strategies to control or alleviate multiple growth-defence trade-offs possibly by perturbing primary metabolic responses. Apart from this, our results demonstrated how metabolic modelling is a useful expansion of transcriptomics analysis and a complement to the analysis of immune signalling network responses ^8,15,18^, as it can highlight potentially different underlying aspects of growth-defence mechanisms that are not clearly observable at the gene regulatory or molecular signalling levels ^81,82^.

To our knowledge, this is presently the most comprehensive metabolic model of potato ^29^, the third most consumed food crop globally. The secondary metabolism reconstruction is also useful across the *Solanaceae* family, which includes staples, such as tomato and pepper, as well as *Nicotiana* species ^83^. These economically important plants have large amounts of published data on biotic as well as abiotic stress and combinations thereof available, and are close relatives of potato, meaning that they share with it the majority of their metabolism, including secondary metabolism ^33,34,84,85^. Therefore, repurposing potato-GEM to study related organisms requires merely (i) adapting the secondary metabolism, (ii) reconfiguring the biomass function and (iii) redefining gene-protein-reaction associations, by finding orthologs from potato or Arabidopsis or obtaining them from online databases. Besides repurposing potato-GEM across related species, there are potentially multiple further avenues of future development with this model, such as multi-tissue ^27,86^ and multi-species ^87,88^ modelling. Moreover, as a further option to increase the capacity of interpreting growth-defence tradeoffs, the metabolic model could be integrated with signalling and regulatory networks, within larger multi-domain modelling frameworks ^81,82^. Apart from cell maintenance and growth, this would enable capturing also the decision-making activities of the cell, such as sensing and responding to environmental changes and regulating metabolism through the activities and abundances of enzymes ^16,89^, likely resulting in a more complete and accurate picture of molecular responses and trade-offs. Therefore, Potato-GEM represents a highly useful resource to study and broaden our understanding of potato as well as general plant defence responses under different types of biotic and abiotic stress.

## 4. Methods

### M1. Metabolic model construction

To develop a reconstruction of potato metabolism, we used three existing genome scale metabolic models (GEMs): (i) model of *Arabidopsis thaliana* core metabolism (AraCore, spanning 549 reactions and 407 metabolites) ^25^, (ii) the basic single-tissue model of tomato metabolism from the Virtual Young TOmato Plant (VYTOP, spanning 2,261 reactions and 2,097 metabolites) ^27^, and (iii) Plant Lipid Module (spanning 5,956 reactions and 3,108 metabolites) ^24^. To enable model intercompatibility and merging, the different reaction and metabolite identifiers of the older AraCore and VYTOP models were updated, including BioCyc ^35^ identifiers, Enzyme Commission (EC) numbers, KEGG ^90^ identifiers, International Chemical Identifier (InChI) strings ^91^ and MetaNetX ^92^ MNXref identifiers, using manual searching and custom Python scripts. Prior to model merging, VYTOP was further curated to remove reactions not supported by present knowledge or located in erroneous compartments.

To merge the AraCore and VYTOP models, each reaction was described as a vector of its metabolites, where each metabolite was represented using its MNXref and compartment identifiers that enable maximum uniqueness. Then, the cosine distance (Python package scipy v1.9.3) ^93^ among the metabolite vectors was used to determine the overlapping reactions between the models that were merged. A cosine distance threshold of 0.2 was used and all non-perfectly overlapping reaction pairs were manually checked to remove spurious merges. Next, the AraCore-VYTOP merged model was extended by integrating a lipid metabolism reconstruction from the Plant Lipid Module ^24^. Prior to this, the merged model was verified according to several criteria, including (i) having a COBRA model structure ^94^; (ii) including metabolites with neutral and/or charged molecular formulas, accompanied by the respective charges and identifiers (e.g., KEGG, ChEBI, PubChem and/or Lipid Maps); and (iii) being able to import and export the model with standard software packages/toolboxes (e.g., via COBRA Toolbox) ^95^. Next, the accompanying software tool that allowed semi-automated integration of the lipid network was used ^24^, which was followed by the removal of reactions that were found to be duplicated.

After merging the three metabolic models, we manually curated the potato secondary metabolism of the resulting model according to the Plant Metabolic Network database v15 (PotatoCyc v6.0.1) ^34,35^. The database was accessed both via its web interface (pmn.plantcyc.org) as well as the BioCyc software Pathway-tools v25.5 ^45,96^ and the PythonCyc v2.02 package. Additionally, the KEGG ^90^ database (/www.genome.jp) and published literature were queried for specific information on phenylpropanoid precursors ^97^, glycoalkaloid ^98,99^ and calystegine metabolism ^100^ as well as cross-compartmental biosynthesis and transports in relation to shikimate ^101^ and hormones biosynthesis ^102–105^. For all secondary pathways the capacity to produce their precursors was verified, where 6 compounds (cytosol L-quinate and D-glucarate for caffeoylglucarate biosynthesis, UDP-D-apiose for apigenin glycosides biosyn., sinapoyl-CoA for hydroxycinnamic acid tyramine amides biosyn., tRNA-Adenosines-37 for cis-zeatin biosyn. and CDP-diacylglycerol for D-myo-inositol(1,4,5)-trisphosphate biosyn.) that currently are not produced by the model due to missing entries in the databases, are fed externally as sinks. In total, 363 reactions were curated, where an additional 277 reactions and 256 metabolites were added to the model. The final secondary metabolic module thus spans 600 reactions from 107 secondary pathways and 22 related precursor pathways and can produce 182 distinct secondary metabolites.

Finally, metabolite chemical formulas and charges were updated according to the MetaNetX resource ^92^ and InChI strings using automated procedures, including custom Python scripts and the Matlab COBRA *getChargeFromInChI* function, as well as manual curation. Custom Matlab scripts were used for full reaction mass and charge balancing, with the exception of 131 reactions that remain charge imbalanced due to chemical formula and charge discrepancies with specific metabolites. The complete procedure resulted in the Potato-GEM model spanning 8,272 reactions and 4,982 metabolites across 16 unique compartments.

### M2. Constraint-based modelling procedures

For constraint-based analysis of metabolic models, the Matlab COBRA v2.42.0 ^95^ and Python COBRApy v0.26.2 ^106^ toolboxes were used. The IBM CPLEX optimization studio v12.10.0 and Gurobi Optimizer v9.5.1 were used with the Matlab and Python toolbox versions, respectively. For the phototrophic (light) regime, the metabolites NO_3_, NH_4_, SO_4_, PO ^3-^, H, H_2_O, CO, Ca, Cl, Fe, K, Mg, Na, H_2_S and photon were set to sink reactions, whereas in the heterotrophic (dark) regime, starch and O_2_ replaced photon and CO_2_ from the phototrophic regime. Note that nitrogen resources refer to the molecules NO_3_ and NH_4_.

To compute how much a unit increase of flux through secondary pathway production (defence) decreases the flux through the biomass reaction (growth) (see Figure 2B), we first computed the flux bounds of each secondary pathway final reaction at the relative growth rate of 0.4 using flux variability analysis (FVA) ^47^. The growth-defence trade-off factor for each secondary pathway was then obtained as the difference between the growth rate (optimal biomass flux obtained using flux balance analysis, FBA) ^74^, obtained by overstepping the maximum secondary metabolite production (FVA upper bound) by one unit, and the growth rate obtained at the maximum secondary metabolite production. The analysis of resource limitation (Figure 3A) was performed by proportionally limiting (within a range of 0 to 1) or expanding (within a range of 1 to 10) resource inputs, including CO_2_, light (photon flux) and nitrogen or a combination of all three, according to the fluxes of these resources obtained at the optimal growth rate of 1 (Figure 2C). Growth rates (optimal biomass fluxes) were then computed at the different ratios of resources using FBA. Lower and upper bounds of flux through secondary pathways were computed using FVA at the secondary metabolism-optimal growth rate of 0.4 to ensure that the largest amount of possible active secondary metabolism was observed.

Randomised Monte Carlo sampling was performed using the Matlab COBRA-implemented Artificial Centering Hit-and-Run (ACHR) algorithm ^67,107^, where 5,000 flux samples per each reaction were computed at 200 steps per point, after 20,000 warmup points to ensure convergence. To realistically constrain the sampling solution space, global upper and lower reaction bounds were computed from maximum and minimum values, respectively, of FVA results obtained at fractions of biomass optimum between 0.4 and 1, in 0.1 incremental steps. Here, with the transcriptomics-constrained models of control conditions, either the final jasmonic acid (CPB) or salicylic acid (PVY) producing reaction upper bounds were set to the tolerance cutoff of 1e-6.

### M3. Experimental procedures

To capture potato biotic stress responses with the Colorado Potato Beetle (CPB), potato cv. Desiree plants were propagated from stem node tissue cultures and transferred to soil 2 weeks after node segmentation. The plants were then kept in growth chambers under controlled environmental conditions at 22/20°C with a long-day (16 h) light photoperiod (light intensity 4,000 lm/m2) and 45–60% relative humidity. For the CPB treatment, two starved 3rd instar larvae were placed on the potato on the first fully developed leaf and left feeding for 30 minutes (Figure 4A). The damaged part of the leaf was excised using a scalpel and immediately frozen in liquid nitrogen. Total RNA was isolated and processed as described in Petek et al. (2014) ^8^. Poly-A selected RNA-seq library prep and paired-end 150 nt Illumina sequencing was performed by Novogene Inc.

To determine the ratio of dry weight to fresh weight, leaves of 22-day old potato cultivar Desiree plants were weighed (fresh mass) and then lyophilized (one day in lyophilizer Christ Gamma 1-20) and weighed again (dry mass). To determine protein content, total proteins were extracted from leaves of the same plants in an extraction buffer containing 10 mM Tris-HCl (pH = 7.5), 150 mM NaCl, 2 mM MgCl and 1 mM DTT (leaves:buffer ratio was 1:2.5). Total protein concentration was estimated using Bio-Rad Protein Assay based on the Bradford dye-binding method ^108^. Measurements were performed using the Synergy MX Multi-Mode Microplate Reader (BioTek). Total protein content was calculated on 1g of leaf tissue.

### M4. Transcriptomics data analysis

To process the transcriptomics data obtained with CPB experiments (see Methods M3), we performed quality control of reads, merging of overlapping pairs, mapping of reads to the potato genome (PGSC v4.03 genome model) ^109^ and read counting using the CLC genomics workbench v12.0 (Qiagen). Differential expression analysis was performed using a standard R package DESeq2 v1.42.1 ^110^ workflow. Genes with a Benjamini Hochberg (BH)-adjusted *p*-value below 0.05 were considered differentially expressed (Figure 4D, Supp. file S5). Gene set enrichment analysis (GSEA) ^60^ was performed using MapMan ontology ^52^ as the source of the gene sets (obtained from GoMapMan database) ^59^ and processed with custom R scripts (Supp. file S6). Principal-coordinate analysis of Bray-Curtis dissimilarity and hierarchical clustering of sample replicates across experiments and treatments was performed using the R packages vegan v2.6-4 ^111^, ggordiplots v.0.4.3 ^112^, and pheatmap v1.0.12 ^113^. Processed transcriptomic data of potato biotic stress responses with Potato Virus Y was obtained from Lukan et al. 2020 ^18^, where the RNA-seq analysis of the tissue within the lesion section A was used as treatment and mock as control.

### M5. Revision of gene-protein-reaction associations

To link the reconstructed model’s reactions with the genes responsible for expressing the underlying enzymatic components (termed gene-protein-reaction, GPR associations) ^64^ we applied the following procedure. Arabidopsis gene identifiers were translated to potato gene identifiers for reactions from AraCore ^25^ and Plant Lipid Module ^24^ with GPRs assigned. Of the 1,014 unique Arabidopsis gene identifiers in total contained within GPRs from these modules, 980 were successfully translated to potato gene identifiers.

The Arabidopsis to potato gene id translation was based on a hierarchical procedure of translations, where initially the PLAZA v5.0 ^61^ comparative genomics resource was used as a source of orthologous gene families in Arabidopsis and potato, serving as the basis to translate Arabidopsis to potato gene identifiers. Of a starting pool of 6,511 Arabidopsis gene identifiers, present in the combined AraCyc v17.1.1 (Plant Metabolic Network v15) database ^34^ as well as AraCore ^25^ and Plant Lipid Module ^24^, we found 6,467 present in PLAZA, 5,872 occurring across 2,122 families with at least one potato gene (and 595 Arabidopsis genes occur in families without any potato gene). We extracted Arabidopsis (Araport11) ^114^, potato (JGI v4.03) ^109^, tomato (ITAG4.0) ^33^, and poplar (JGI v4.1) ^115^ gene sequences from the families with a sufficient number of genes (1,708 families), and performed all to all computations of sequence Levenshtein distances (R package stringdist v.0.9.10) ^116^ between the genes within each family. Three of the families had high similarity between all genes (distances of less than 0.08), and we assumed all-to-all translations for these. For the others, we transformed the Levenstein distance to a similarity score (using e^(–x^3), scaled between 0 and 1), and applied a “whole-graph” similarity threshold per family (the maximum threshold whereby each gene still has at least one similarity to another gene above the threshold) ^117^. For the families with 100 or fewer members, we performed Louvain clustering (Python package louvain v0.8.2, incrementally increasing the threshold if clustering did not converge) ^118^, resulting in translation groups for 3,347 of the Arabidopsis genes. For the remaining 32 families with more than 100 members (too large to cluster using Louvain) we performed Markov clustering (inflation parameter of 5 used) ^119^, resulting in translation groups for a further 624 genes. Each of these translation groups contained at least one Arabidopsis and at least one potato identifier for 3,975 Arabidopsis genes, which rendered translations to potato for 712 of the 1,014 Arabidopsis GPR gene identifiers.

Next, the remaining gene identifiers were translated using BLAST+ v2 ^62,120^ reciprocal best hit search on Arabidopsis vs. potato PGSCv4 proteomes, where the highest scoring hits were used as translations. Default parameters were used with both approaches, with the top three potato gene identifiers per Arabidopsis identifier considered. ITAG potato identifiers were additionally consolidated to PGSC using an *in-house* generated translation table (files available from https://github.com/NIB-SI/DiNAR/tree/master/TranslationTables) ^121^. In case of many-to-many relationships, the translation pair was prioritised based on the differential expression and raw count values, selecting the most up-or down-regulated gene.

Finally, the model was revised with potato-translated GPR associations by preferentially including translated GPRs from AraCore and then from the Plant Lipid Module, for 315 and 3036 reactions, respectively, and finally GPRs from the PotatoCyc v6.0.1 (Plant Metabolic Network v15) database ^34^ for a further 806 reactions.

### M6. Statistical data analysis

For statistical hypothesis testing, Scipy ^93^ v1.9.3 and Statsmodels v0.14.0 were used with default settings. All statistical tests were two-tailed except where stated otherwise. For correlation analysis, the Spearman correlation coefficient is reported and statistical significance was tested using Welch’s t-test. To correct for multiple comparisons and control the false discovery rate, the Benjamini-Hochberg method was used.

For differential flux analysis of reaction flux samples obtained by performing randomized Monte Carlo sampling (see Methods M2), results across transcriptome-constrained models of experiment treatments (triplicates) were pooled and reaction tolerance cutoff of 1e-6 was used. The Kolmogorov Smirnov test was used to assess the significance of differential flux distributions, and log2 fold-change scores of median values of the flux samples between treatment and control conditions were computed per reaction. Reactions with significant 2-fold or higher differences in flux values between treatment and control conditions were regarded as exhibiting differential fluxes. Afterward, pathway enrichment was performed to determine pathways that are enriched in these differential flux-exhibiting reactions using Fisher’s exact test based on the hypergeometric distribution.

To analyse the correlations between the sampled fluxes of the enriched secondary and primary pathways, absolute flux values were summed over the respective pathway reactions to obtain the total pathway flux and the direction of correlation was determined according to the direction of regulation (up/down) of the pathways in question. The pairwise correlation profiles were clustered using the cosine distance to characterize the similarity of two correlation vectors. Permutational tests were performed to evaluate the significance of the similarity according to the Weibull distribution, where the null similarity scores were defined by randomizing the first vector 10,000 times and recomputing the similarity between the randomized first and non-randomized second vectors.

### M7. Software and data

Python v3.8.16 (www.python.org), R v4.3.1 (www.r-project.org/) and Matlab v2019b (www.mathworks.com) were used for computations. For general data analysis and visualisation, the Python packages Pandas v1.5.2, Biopython v1.78, and Seaborn v0.13.2 were used. The model and related files, including Supp. files S1-S6, as well as code to reproduce the results were deposited to the Github repository and are available at https://github.com/NIB-SI/Potato-GEM. Source data was deposited to the Zenodo repository at https://doi.org/10.5281/zenodo.xxxxxxx.

## Acknowledgements

We thank Phillip Wendering and Anže Županič for technical discussions and critical comments, as well as Živa Ramšak for technical support. The study was supported by the Slovenian Research and Innovation Agency (ARIS) grants no. J2-3060 (J.Z.), Z4-50146 (C.B.) and P4-0165 (K.G.), the Public Scholarship, Development, Disability and Maintenance Fund of the Republic of Slovenia grant no. 11013-9/2021-2 (J.Z.), the European Union Horizon 2020 Framework Programme under grant agreement no. 862858 (https://adapt.univie.ac.at/, K.G.). The European Union’s Horizon 2020 research and innovation program also financially supported this work through grant 862201 (to Z.N.). Z.N and S.C.C. acknowledge the financial support by the Collaborative Research Center 1644 funded by the German Research Foundation.

