## Supplementary information for "Evaluating plant growth-defence trade-offs by modelling the interaction between primary and secondary metabolism"

† shared first authorship

#### Table of contents

Supplementary Figures … p.2 - p.20
Supplementary Tables … p.21 - p.25
Supplementary References … p.26

### Supplementary figures

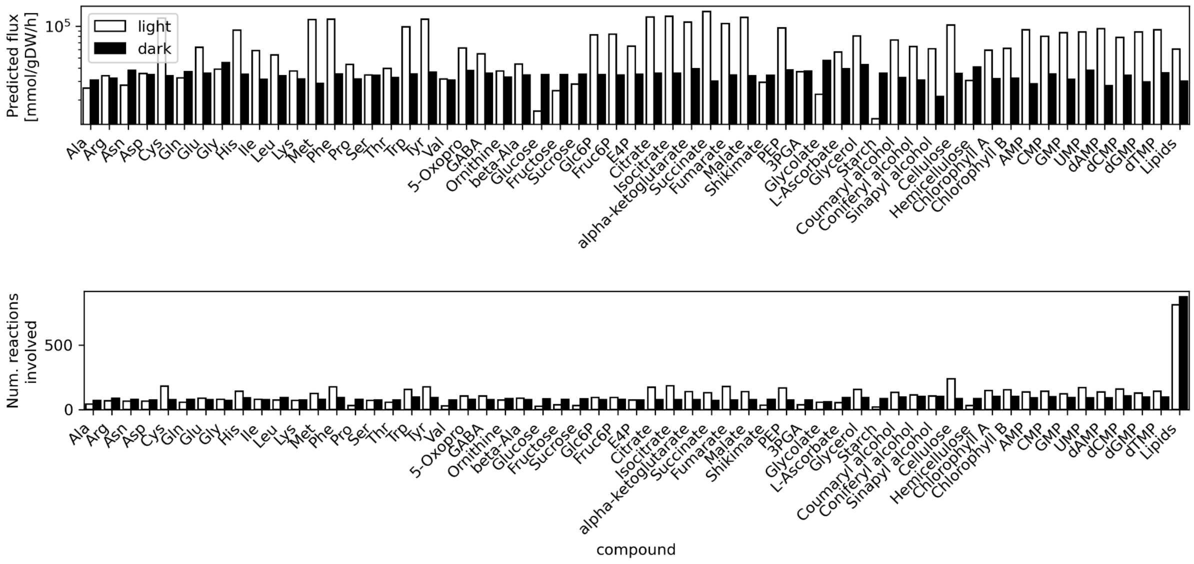


**Figure S1.1** Biomass component production.

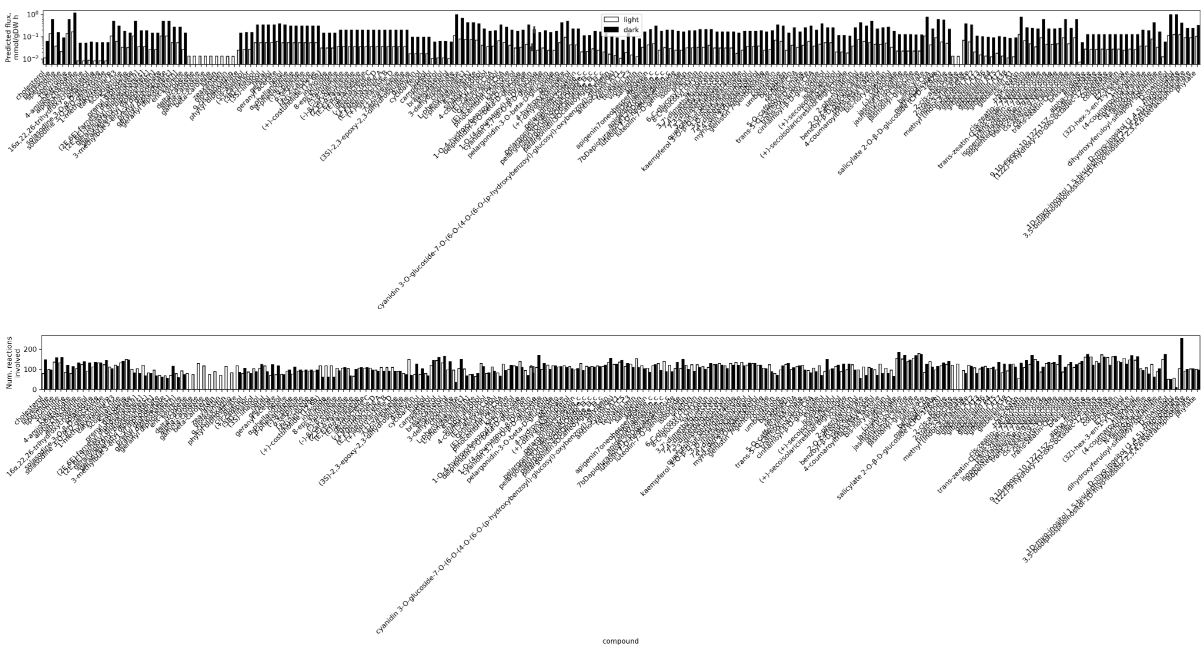


**Figure S1.2** Secondary metabolite production.

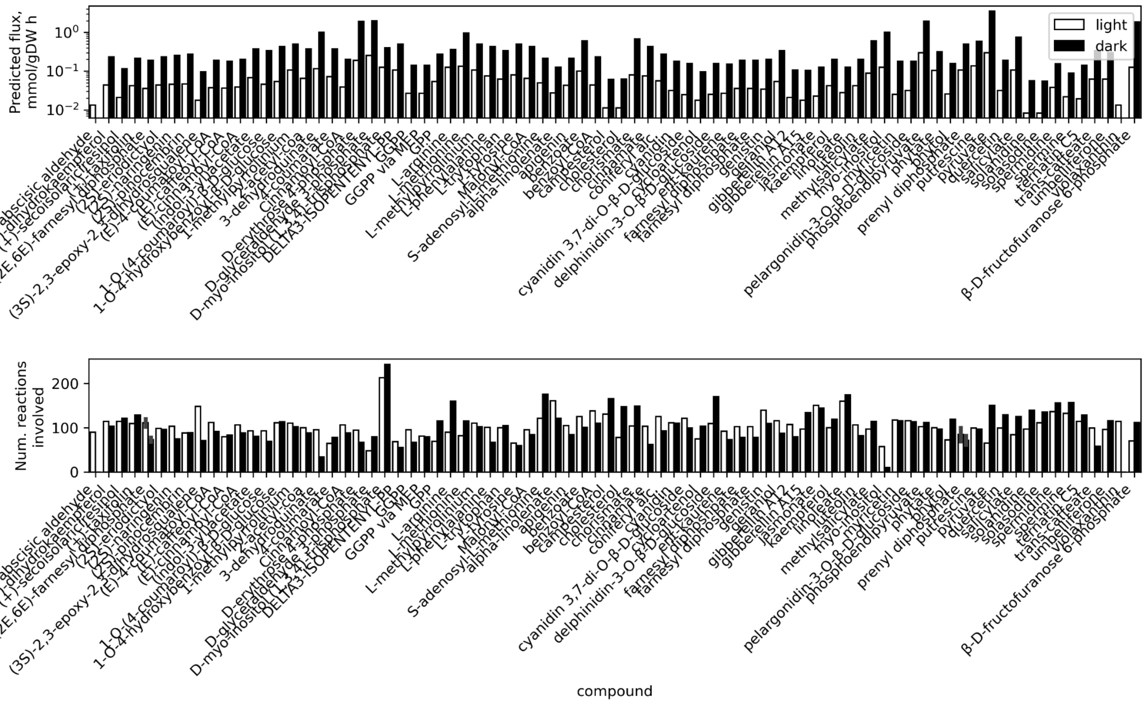


**Figure S1.3** Production of secondary metabolism-related precursors.

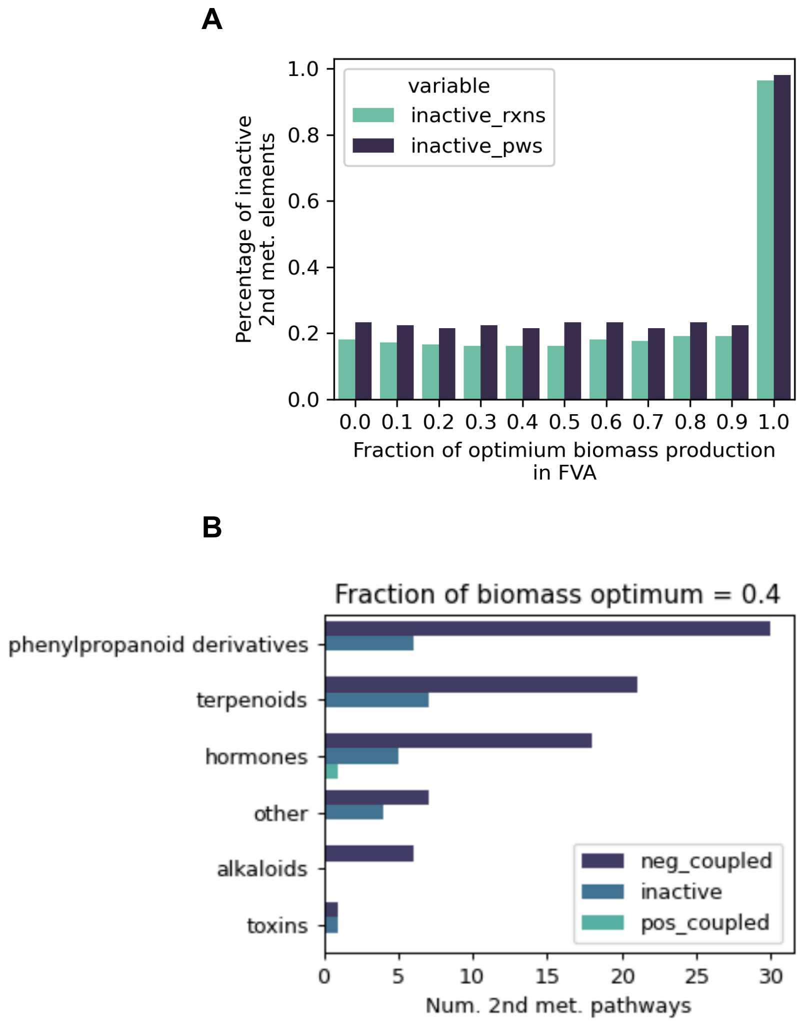


**Figure S2.1** (A) Ratio of inactive secondary pathways and reactions in response to the fraction of optimum biomass production set in FVA. (B) Number of active and inactive secondary pathways at a relative growth rate of 0.4.

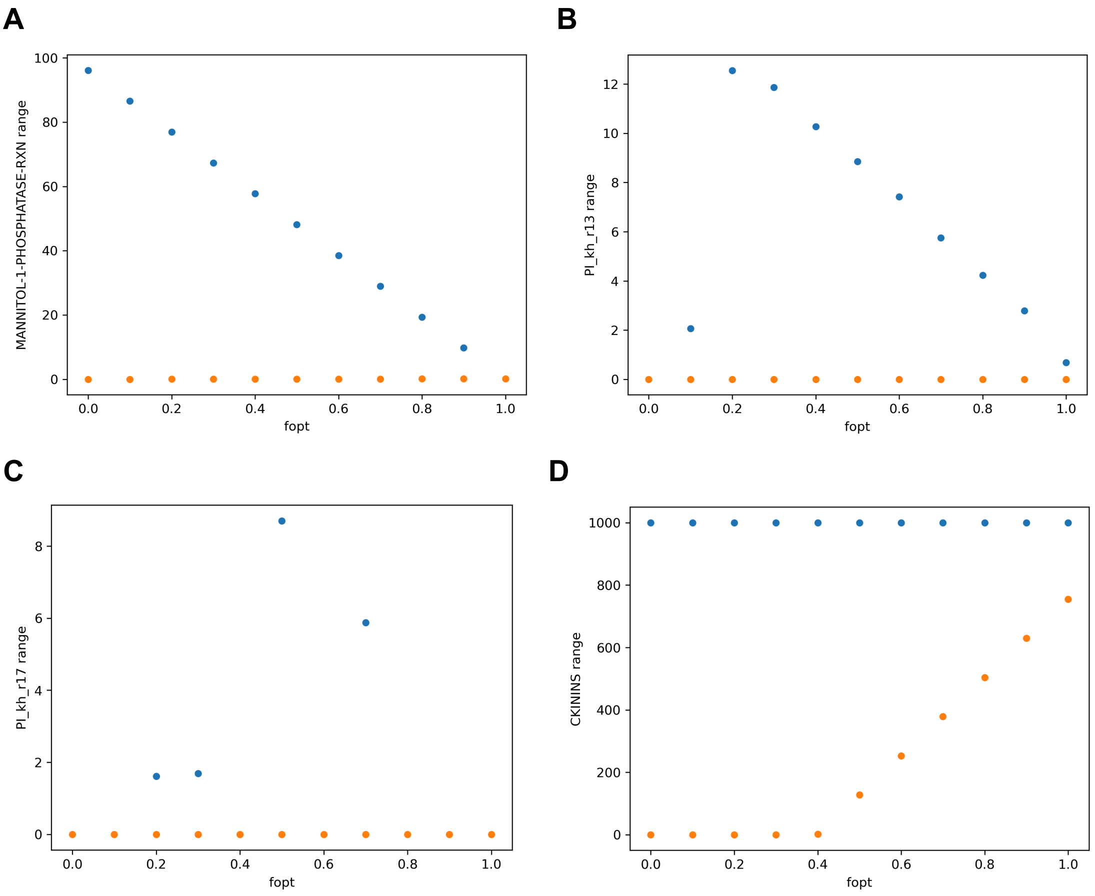


**Figure S2.2** Demonstration of the different types of observed coupling between the potato basic growth and stress response modes. The variability of fluxes of the final reactions in different secondary pathways were evaluated within the optimal biomass space using flux variability analysis (FVA) [1] across different fractions, ranging from 0 to 1, of the optimal relative growth rate were used (denoted 'fopt' on the x-axis). (A, B) negative coupling, where (B) is non-negative at a relative growth rate of 1 and decreases to zero at the other growth rate extreme. (C) Unstable. (D) Positive coupling, where the lower bound increases with an increasing relative growth rate (cytokinin biosynthesis, PWY-2681).

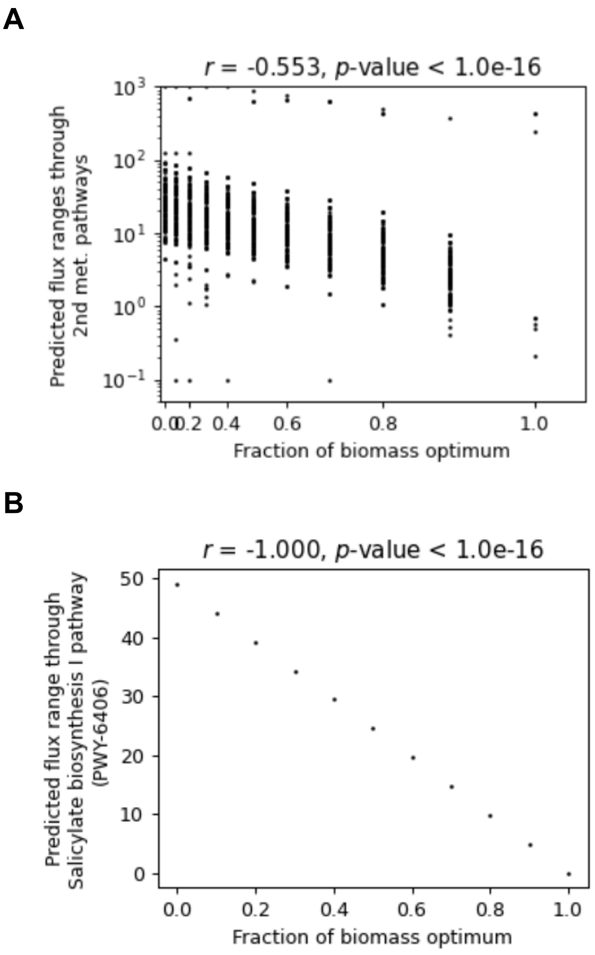


**Figure S2.3** Spearman correlation analysis between secondary metabolite production and growth (fraction of biomass optimum), with (A) absolute flux ranges across all secondary pathways, and (B) absolute flux range of an exemplary negatively coupled pathway, Salicylic acid biosynthesis (PWY-6406).

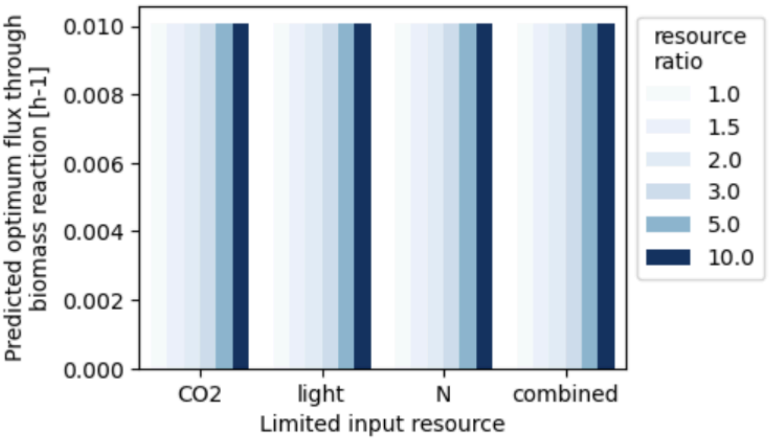


**Figure S3.1** Effect of resource expansion, at a ratio of 1, 1.5, 2, 3, 5 and 10, on the predicted relative growth rate (biomass optimum).

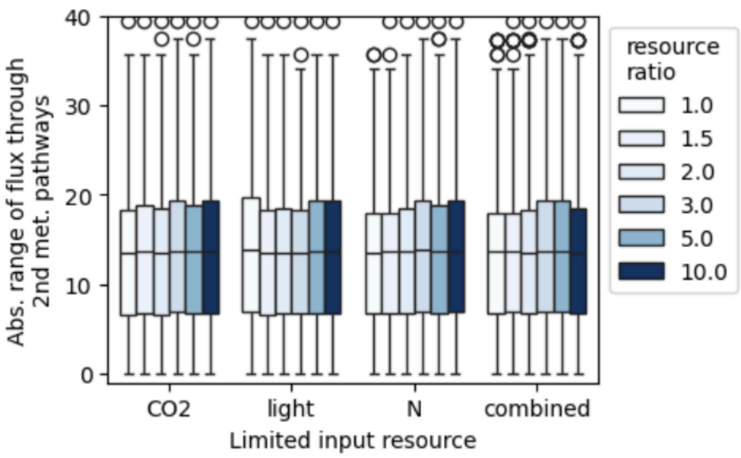


**Figure S3.2** Effect of resource expansion, at a ratio of 1, 1.5, 2, 3, 5 and 10, on the range of secondary metabolite production (defence), computed at a fraction of 0.4 of the relative growth rate to observe the largest amount of possible active secondary metabolism.

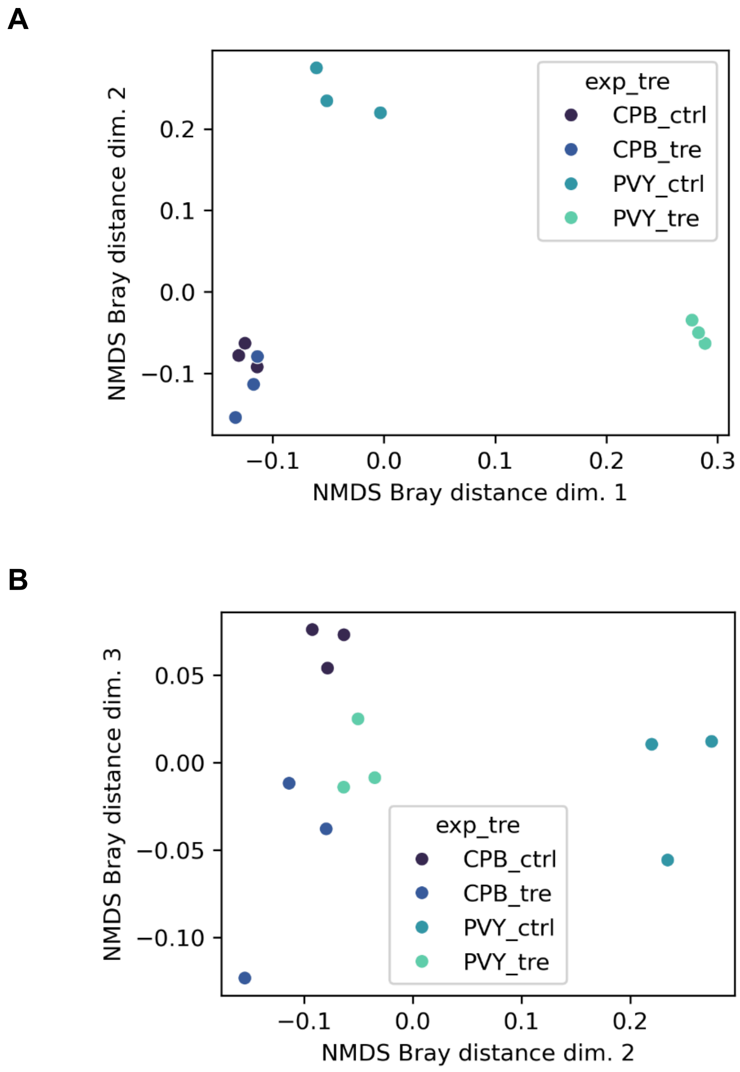


**Figure S4.1** Principal-coordinate analysis of Bray-Curtis dissimilarity between transcript counts across experiments and treatments, showing (A) dim. 1 vs. dim. 2 and (B) dim. 2 vs. dim. 3.

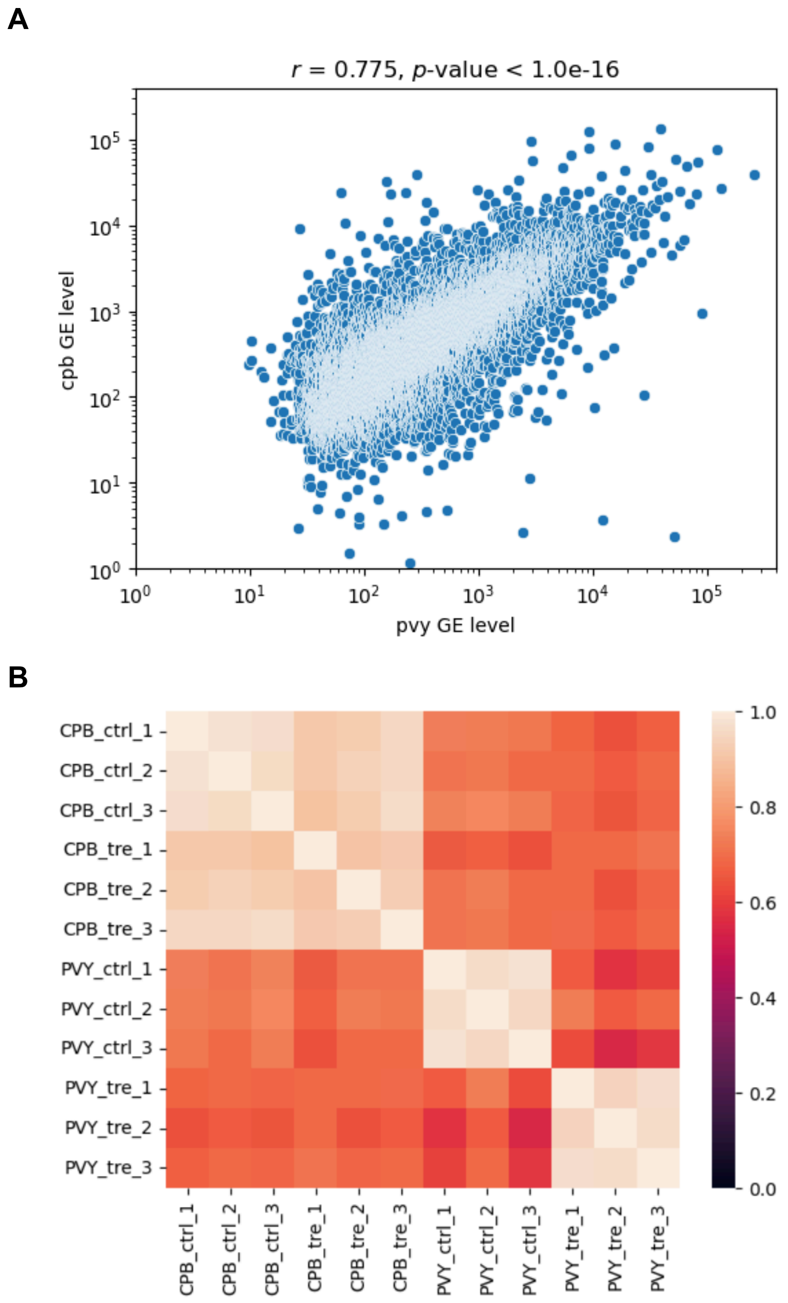


**Figure S4.2** Spearman correlation analysis. (A) Correlation scatter plot of the average of transcriptomics experiment controls among CPB and PVY experiments. (B) Heatmap of correlations among all replicates across treatments (treatment: tre, control: ctrl) and experiments (CPB, PVY).

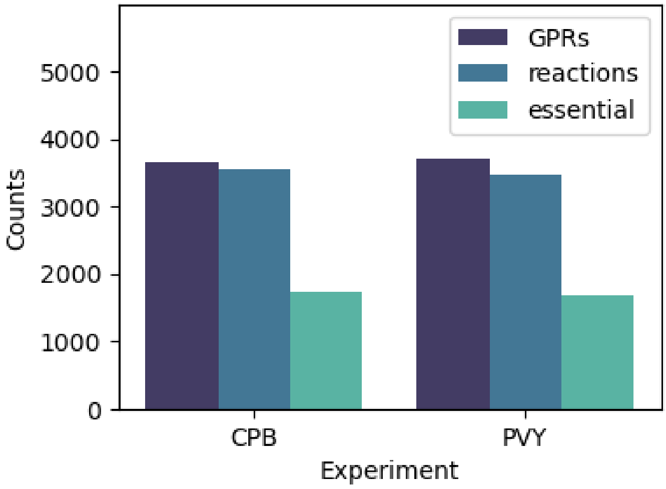


**Figure S5.1** Comparison of the number of unique protein complexes in GPR rules (denoted GPRs), GPR-annotated reactions (reactions) and essential reactions (essential) in the CPB and PVY transcriptome-constrained models. 14,404 and 19,367 transcripts from the CPB or PVY experiment, respectively, were were mapped to 3,662 and 3,716 unique protein complexes in GPR rules across 3,561 and 3,467 reactions, spanning 1,725 and 1,692 essential reactions, respectively.

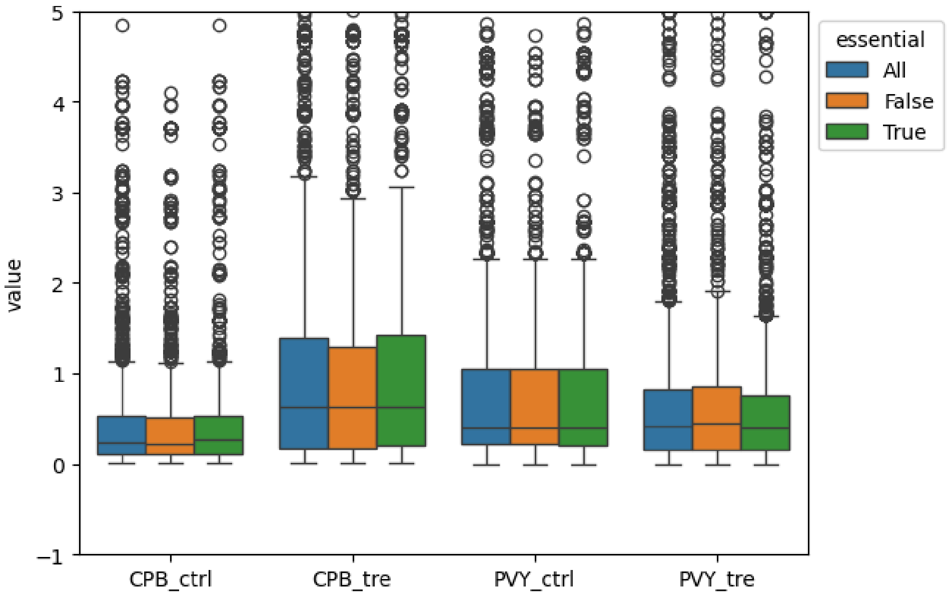


**Figure S5.2** Variability of transcriptome-constrained model's upper bounds with all, essential and non-essential reactions across experiments and treatments.

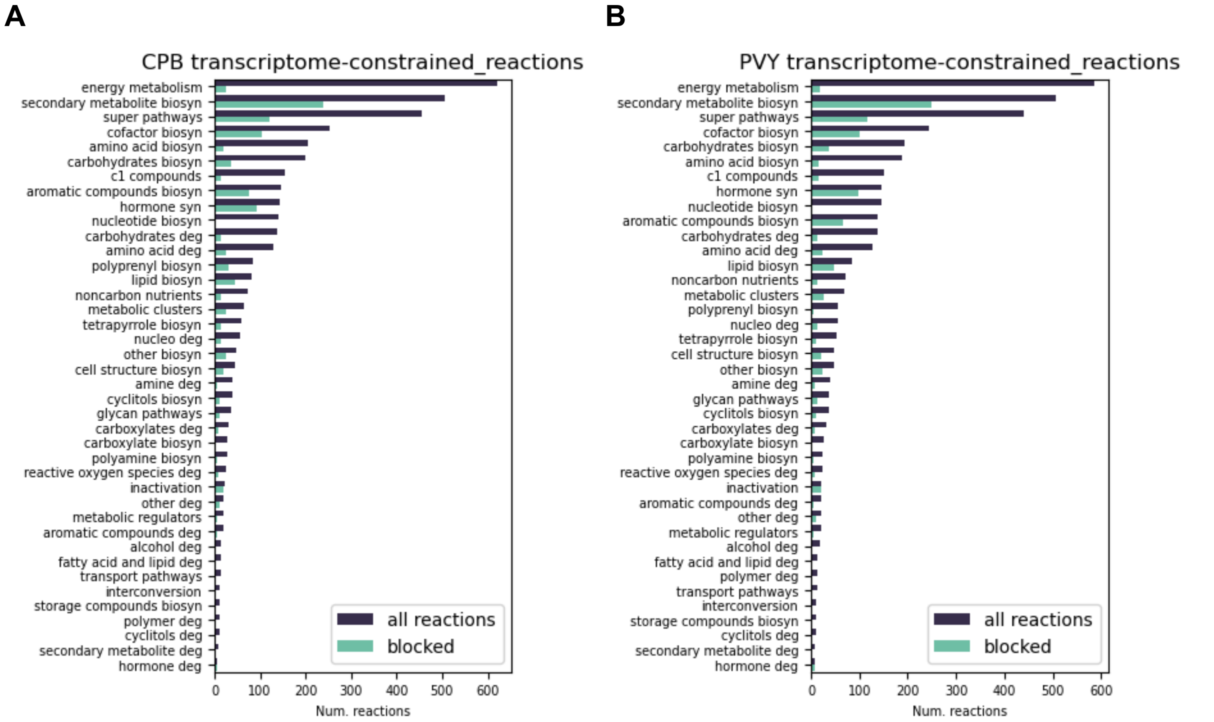


**Figure S5.3**. Pathway ontologies across (A) CPB and (B) PVY transcriptome-constrained models.

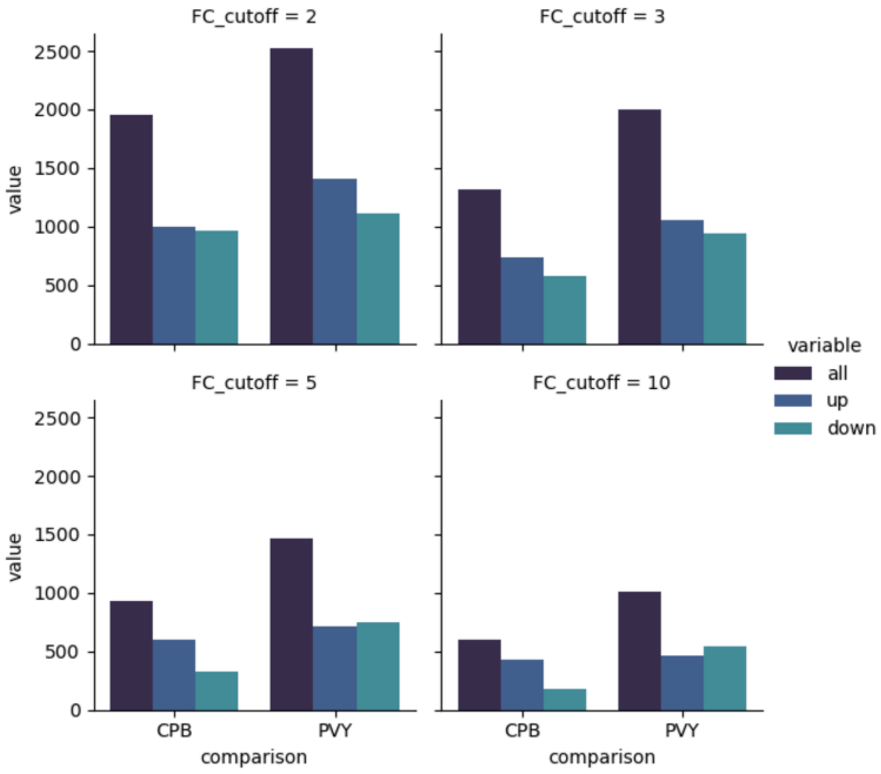


**Figure S6.1** Number of identified reactions with differential fluxes at different fold-change cutoffs ranging from 2 to 10 (Kolmogorov Smirnov test BH-corrected $p$-value < 0.05). The number of upregulated (up), downregulated (down) and total (all) reactions are depicted.

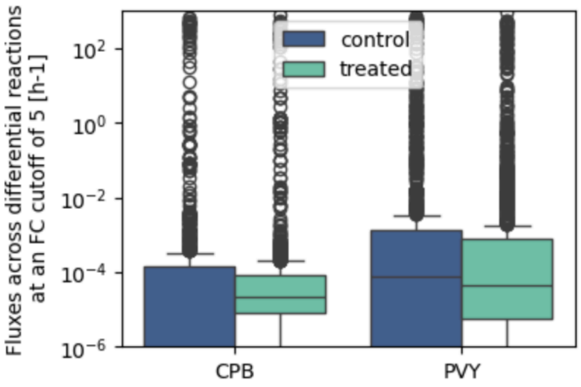


**Figure S6.2** Fluxes of the identified differential reactions at a fold-change cutoff of 5 (Kolmogorov Smirnov test BH-corrected $p$-value < 0.05) in treated and control samples of CPB and PVY experiments.

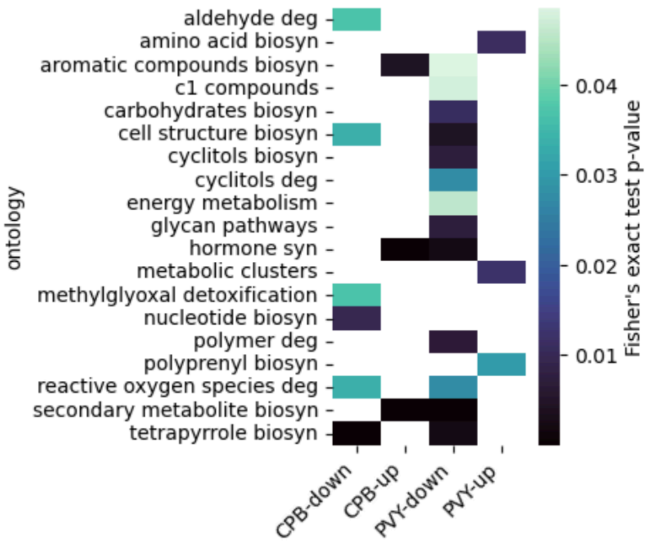


**Figure S6.3** Enrichment (Fisher's exact test BH-corrected p-value < 0.05) of metabolic subsystems at the BioCyc [2] ontology level.

**A**

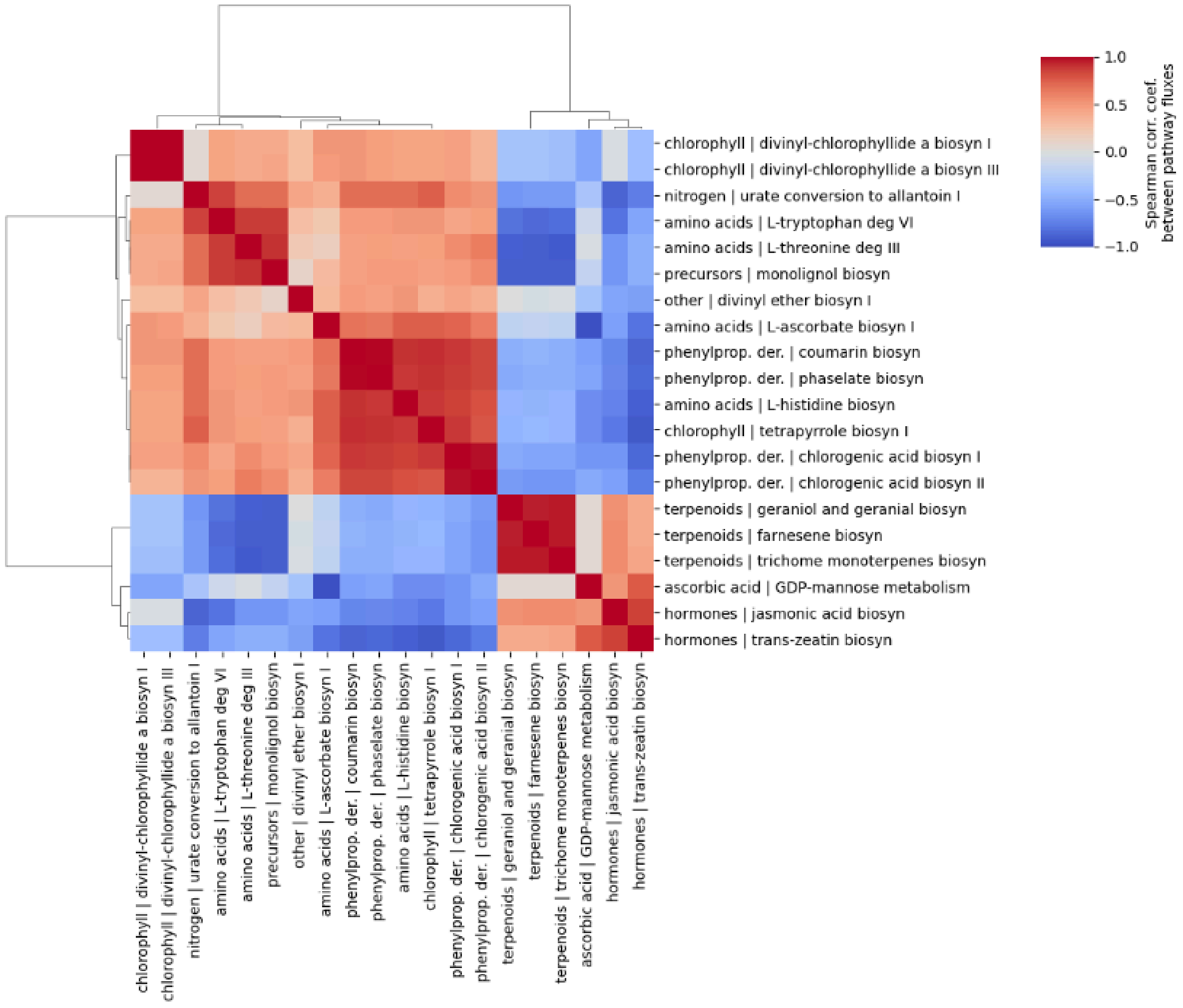
chlorophyll | divinyl-chlorophyllide a biosyn I
chlorophyll | divinyl-chlorophyllide a biosyn III
nitrogen | urate conversion to allantoin I
amino acids | L-tryptophan deg VI
amino acids | L-threonine deg III
precursors | monolignol biosyn
other | divinyl ether biosyn I
amino acids | L-ascorbate biosyn I
phenylprop. der. | coumarin biosyn
phenylprop. der. | phaselate biosyn
amino acids | L-histidine biosyn
chlorophyll | tetrapyrrole biosyn I
phenylprop. der. | chlorogenic acid biosyn I
phenylprop. der. | chlorogenic acid biosyn II
terpenoids | geraniol and geranial biosyn
terpenoids | farnesene biosyn
terpenoids | trichome monoterpenes biosyn
ascorbic acid | GDP-mannose metabolism
hormones | jasmonic acid biosyn
hormones | trans-zeatin biosyn
1.0
0.5
0.0
−0.5
−1.0
Spearman corr. coef.
between pathway fluxes

**B**

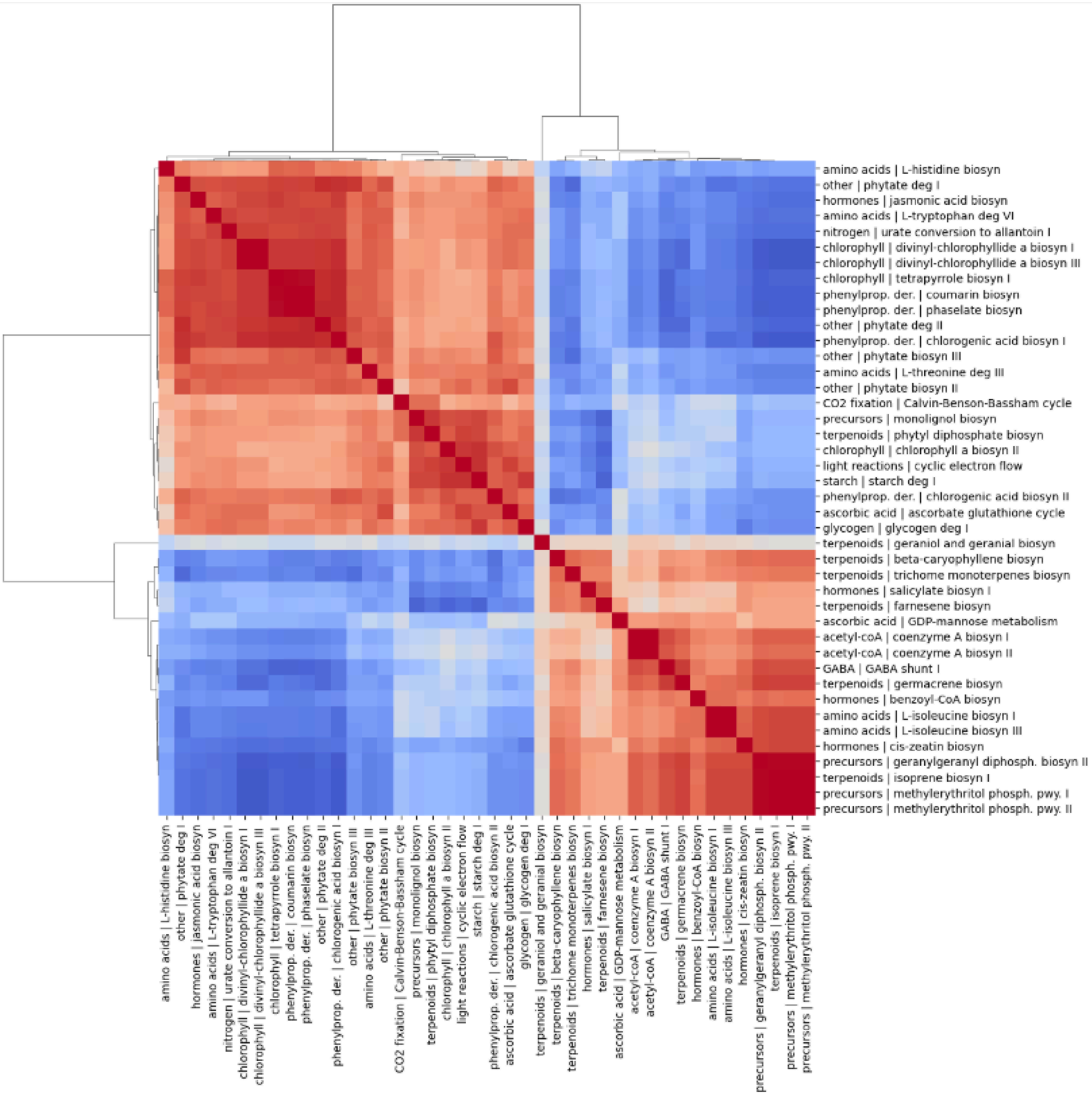


**Figure S6.4** Depiction of (A) CPB and (B) PVY correlations between total fluxes of all enriched (Fisher's exact test BH-corrected *p*-value < 0.05) pathways. Cosine distance used as distance metric and Spearman correlation coefficient shown (n samples = 1,000). Correlations above 0.063 or below -0.063 are significant below the 0.05 significance threshold.

**A**

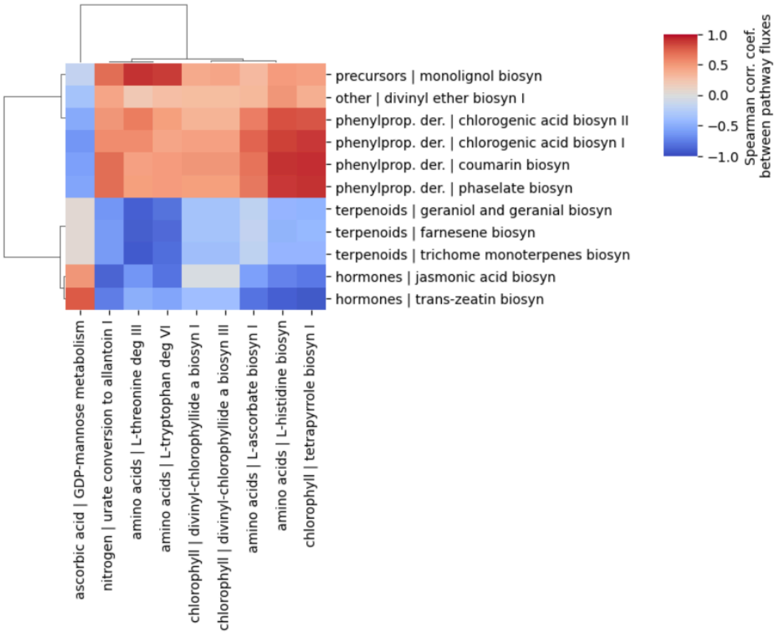


**B**

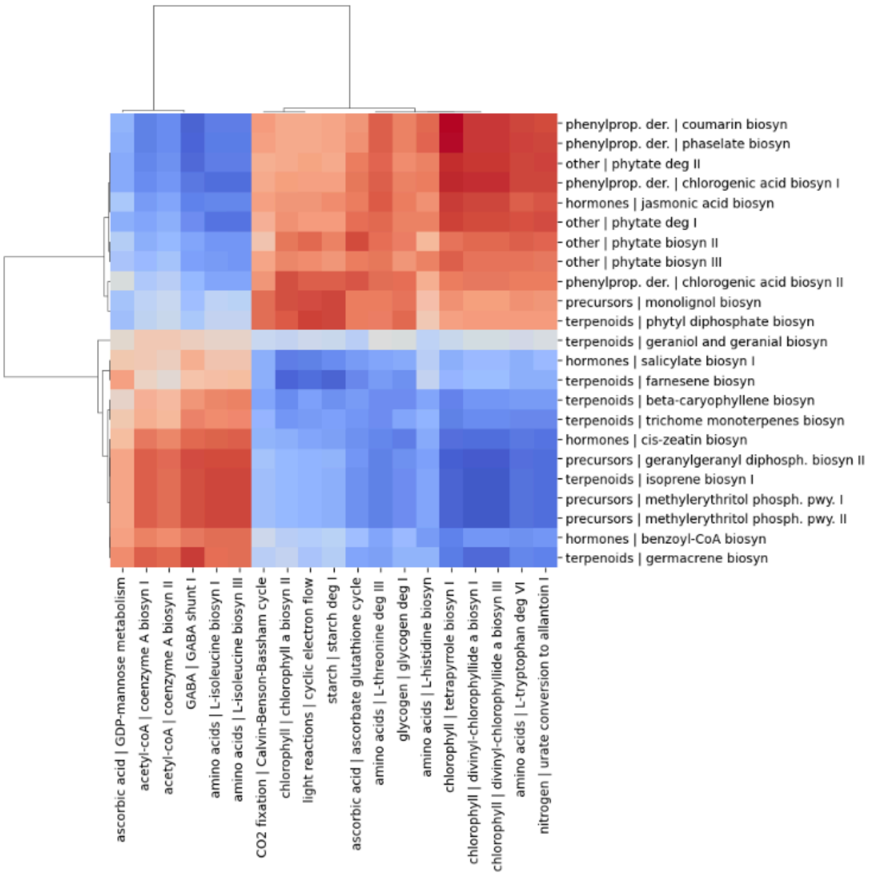


**Figure S6.5** Depiction of (A) CPB and (B) PVY growth-defence tradeoffs as correlations between total fluxes of the enriched (Fisher's exact test BH-corrected *p*-value $< 0.05$) secondary vs. primary pathways. Cosine distance used as distance metric and Spearman correlation coefficient shown (n samples = 1,000).

#### Supplementary tables

**Table S1** Overview of leaf biomass components and sources.

| Compound | g/g DW | mmol/g DW | Source | Organism |
|---|---|---|---|---|
| Proteins | 0.150 | 1.186 | This study | *Solanum tuberosum* |
| Free amino acids | 0.017 | 0.130 | [3–5] | *Solanum lycopersicum, Solanum tuberosum* |
| Sugars and organic acids | 0.265 | 1.436 | [3,4,6,7] | *Solanum tuberosum, Solanum lycopersicum* |
| Lipids | 0.059 | 0.161 | [8] | *Solanum tuberosum* |
| Lignin and (hemi)cellulose precursors | 0.176 | 1.040 | [9] | *Solanum lycopersicum* |
| Chlorophylls | 0.030 | 0.034 | [9] | *Solanum lycopersicum* |
| Minerals | 0.158 | 3.746 | [10,11] | *Solanum tuberosum* |
| Nucleic acids | 0.145 | 0.430 | This study: inferred | *Solanum tuberosum* |

**Table S2** Overview of secondary metabolism pathways and products across classes and subclasses.

| Class | Subclass | Num. pathways | Num. reactions | Num. products |
|---|---|---|---|---|
| **alkaloids** | **alkaloids** | 6 | 41 | 10 |
| **hormones** | **abscisic acid** | 2 | 12 | 3 |
| | **auxins** | 2 | 4 | 2 |
| | **brassinosteroids** | 2 | 46 | 2 |
| | **cytokinins** | 5 | 38 | 11 |
| | **ethanol** | 1 | 3 | 1 |
| | **gibberellins** | 4 | 25 | 9 |
| | **jasmonic acid** | 2 | 28 | 5 |
| | **salicylic acid** | 6 | 24 | 6 |
| **other** | **fatty acid derivatives** | 3 | 19 | 8 |
| | **nitrogen-containing** | 2 | 12 | 5 |
| | **sugar alcohols** | 6 | 38 | 7 |
| **phenylpropanoid derivatives** | **cinnamates** | 5 | 17 | 7 |
| | **coumarins** | 4 | 16 | 4 |
| | **flavonoids** | 20 | 95 | 39 |
| | **lignan** | 2 | 13 | 3 |
| | **lignin** | 1 | 2 | 1 |
| | **other** | 3 | 9 | 3 |
| | **phenylprop. precursors** | 1 | 23 | 10 |
| **terpenoids** | **diterpenoid** | 3 | 7 | 3 |
| | **monoterpenoid** | 7 | 23 | 13 |
| | **sesquiterpenoid** | 5 | 24 | 16 |
| | **tetraterpenoid** | 7 | 24 | 9 |
| | **triterpenoid** | 6 | 30 | 9 |
| **toxins** | **phytoalexins** | 2 | 9 | 4 |

**Table S3** Enrichment of metabolic subsystems (Fisher's exact test BH-corrected p-value < 0.05) at the level of BioCyc ontologies [2], across experiments and conditions (data visualized in Figure S6.3).

| | CPB | CPB | PVY | PVY |
|---|---|---|---|---|
| Ontology | down | up | down | up |
| aldehyde deg | 3.69E-02 | | | |
| amino acid biosyn | | | | 1.12E-02 |
| aromatic compounds biosyn | | 4.00E-03 | 4.85E-02 | |
| c1 compounds | | | 4.75E-02 | |
| carbohydrates biosyn | | | 1.09E-02 | |
| cell structure biosyn | 3.36E-02 | | 4.08E-03 | |
| cyclitols biosyn | | | 7.57E-03 | |
| cyclitols deg | | | 2.77E-02 | |
| energy metabolism | | | 4.58E-02 | |
| glycan pathways | | | 7.57E-03 | |
| hormone syn | | 4.00E-10 | 2.16E-03 | |
| metabolic clusters | | | | 1.22E-02 |
| methylglyoxal detoxification | 3.69E-02 | | | |
| nucleotide biosyn | 9.81E-03 | | | |
| polymer deg | | | 6.61E-03 | |
| polyprenyl biosyn | | | | 3.03E-02 |
| reactive oxygen species deg | 3.36E-02 | | 2.77E-02 | |
| secondary metabolite biosyn | | 1.16E-06 | 2.53E-04 | |
| tetrapyrrole biosyn | 1.85E-04 | | 2.16E-03 | |

**Table S4** Enrichment of metabolic subsystems (Fisher's exact test BH-corrected p-value < 0.05) at the level of BioCyc pathways [2], across experiments and conditions. The data is visualized in Figure 6D,E.

| | | | CPB | CPB | PVY | PVY |
|---|---|---|---|---|---|---|
| Metabolism | BioCyc id | Name | down | up | down | up |
| 2nd - terpenoids | PWY-6447 | trichome monoterpenes biosynthesis | | 7.61E-03 | | 7.22E-06 |
| 2nd - terpenoids | PWY-6275 | beta-caryophyllene biosynthesis | | | | 4.07E-02 |
| 2nd - terpenoids | PWY-6270 | isoprene biosynthesis I | | | | 1.46E-03 |
| 2nd - terpenoids | PWY-5829 | geraniol and geranial biosynthesis | | 7.61E-03 | | 4.07E-02 |
| 2nd - terpenoids | PWY-5733 | germacrene biosynthesis | | | | 4.07E-02 |
| 2nd - terpenoids | PWY-5725 | farnesene biosynthesis | | 7.61E-03 | | 4.07E-02 |
| 2nd - terpenoids | PWY-5063 | phytyl diphosphate biosynthesis | | | 4.79E-02 | |
| 2nd - precursors | PWY-7560 | methylerythritol phosphate pathway II | | | | 4.93E-04 |
| 2nd - precursors | PWY-5121 | geranylgeranyl diphosphate biosynthesis II (via MEP) | | | | 2.09E-03 |
| 2nd - precursors | PWY-361 | monolignol biosynthesis | 5.16E-11 | | 1.38E-04 | |
| 2nd - precursors | NONMEVIPP-PWY | methylerythritol phosphate pathway I | | | | 4.93E-04 |
| 2nd - phenylpropanoid derivatives | PWY-6320 | phaselate biosynthesis | 7.24E-03 | | 1.25E-02 | |
| 2nd - phenylpropanoid derivatives | PWY-6040 | chlorogenic acid biosynthesis II | 1.28E-02 | | 3.78E-02 | |
| 2nd - phenylpropanoid derivatives | PWY-6039 | chlorogenic acid biosynthesis I | 4.91E-02 | | 2.26E-04 | |
| 2nd - phenylpropanoid derivatives | PWY-5868 | coumarin biosynthesis | 1.28E-02 | | 1.25E-02 | |
| 2nd - other | PWY-6362 | 1D-myo-inositol hexakisphosphate biosynthesis II | | | 1.60E-02 | |
| 2nd - other | PWY-5406 | divinyl ether biosynthesis I | 1.28E-02 | | | |
| 2nd - other | PWY-4781 | phytate degradation II | | | 7.46E-03 | |
| 2nd - other | PWY-4702 | phytate degradation I | | | 2.18E-02 | |
| 2nd - other | PWY-4661 | 1D-myo-inositol hexakisphosphate biosynthesis III | | | 3.23E-02 | |

| | | | | | | |
|---|---|---|---|---|---|---|
| 2nd - hormones | PWY-735 | jasmonic acid biosynthesis | | 7.16E-27 | 1.27E-18 | |
| 2nd - hormones | PWY-6458 | benzoyl-CoA biosynthesis | | | | 4.07E-02 |
| 2nd - hormones | PWY-6406 | salicylate biosynthesis I | | | | 4.07E-02 |
| 2nd - hormones | PWY-2781 | cis-zeatin biosynthesis | | | | 5.43E-03 |
| 2nd - hormones | PWY-2681 | trans-zeatin biosynthesis | | 2.95E-03 | | |
| primary | THRDLCTCAT-PWY | L-threonine degradation III (to methylglyoxal) | 8.12E-03 | | 4.79E-02 | |
| primary | PWY-882 | L-ascorbate biosynthesis I (L-galactose pathway) | 4.12E-02 | | | |
| primary | PWY-842 | starch degradation I | | | 1.60E-02 | |
| primary | PWY-8270 | cyclic electron flow | | | 4.79E-02 | |
| primary | PWY-7851 | coenzyme A biosynthesis II (eukaryotic) | | | | 4.07E-02 |
| primary | PWY-7159 | 3,8-divinyl-chlorophyllide a biosynthesis III (aerobic, light independent) | 3.52E-02 | | 5.17E-04 | |
| primary | PWY-5691 | urate conversion to allantoin I | 3.52E-02 | | 4.79E-02 | |
| primary | PWY-5188 | tetrapyrrole biosynthesis I (from glutamate) | 3.52E-02 | | 4.82E-04 | |
| primary | PWY-5103 | L-isoleucine biosynthesis III | | | | 2.24E-02 |
| primary | PWY-5064 | chlorophyll a biosynthesis II | | | 4.41E-02 | |
| primary | PWY-3181 | L-tryptophan degradation VI (via tryptamine) | 8.12E-03 | | 4.79E-02 | |
| primary | PWY-2261 | ascorbate glutathione cycle | | | 2.20E-02 | |
| primary | MANGDPMET-PWY | GDP-mannose metabolism | | 8.60E-03 | | 1.90E-02 |
| primary | ILEUSYN-PWY | L-isoleucine biosynthesis I (from threonine) | | | | 1.90E-02 |
| primary | HISTSYN-PWY | L-histidine biosynthesis | 2.87E-08 | | 6.50E-07 | |
| primary | GLYCOCAT-PWY | glycogen degradation I | | | 4.72E-02 | |
| primary | GLUDEG-I-PWY | GABA shunt I | | | | 4.07E-02 |
| primary | COA-PWY | coenzyme A biosynthesis I | | | | 8.62E-04 |
| primary | CHLOROPHYLL-SYN | 3,8-divinyl-chlorophyllide a biosynthesis I (aerobic, light-dependent) | 6.05E-03 | | 7.62E-05 | |
| primary | CALVIN-PWY | Calvin-Benson-Bassham cycle | | | 2.86E-04 | |
